# Quantifying and Mitigating Motor Phenotypes Induced by Antisense Oligonucleotides in the Central Nervous System

**DOI:** 10.1101/2021.02.14.431096

**Authors:** Michael P. Moazami, Julia M. Rembetsy-Brown, Feng Wang, Pranathi Meda Krishnamurthy, Alexandra Weiss, Miklos Marosfoi, Robert M. King, Mona Motwani, Heather Gray-Edwards, Katherine A. Fitzgerald, Robert H. Brown, Jonathan K. Watts

**Affiliations:** RNA Therapeutics Institute, University of Massachusetts Medical School, Worcester, MA, 01605 USA; Department of Neurology, University of Massachusetts Medical School, Worcester, MA, 01605 USA; Department of Radiology, UMass Medical School, Worcester, MA, 01605 USA; Department of Biomedical Engineering, Worcester Polytechnic Institute, Worcester, MA, 01609, USA; Program in Innate Immunity, Division of Infectious Diseases and Immunology, Department of Medicine, University of Massachusetts Medical School, Worcester, MA, 01605 USA; Department of Biochemistry and Molecular Pharmacology, University of Massachusetts Medical School, Worcester, MA, 01605 USA

**Author notes:** These authors contributed equally to the work (Michael P. Moazami, Julia M. Rembetsy-Brown).

## Abstract

Antisense oligonucleotides (ASOs) are emerging as a promising class of therapeutics for neurological diseases. When injected directly into the cerebrospinal fluid, ASOs distribute broadly across brain regions and exert long-lasting therapeutic effects. However, many phosphorothioate (PS)-modified gapmer ASOs show transient motor phenotypes when injected into the cerebrospinal fluid, ranging from reduced motor activity to ataxia or acute seizure-like phenotypes. The effect of sugar and phosphate modifications on these phenotypes has not previously been systematically studied. Using a behavioral scoring assay customized to reflect the timing and nature of these effects, we show that both sugar and phosphate modifications influence acute motor phenotypes. Among sugar analogues, PS-DNA induces the strongest motor phenotype while 2’-substituted RNA modifications improve the tolerability of PS-ASOs. This helps explain why gapmer ASOs have been more challenging to develop clinically relative to steric blocker ASOs, which have a reduced tendency to induce these effects. Reducing the PS content of gapmer ASOs, which contain a stretch of PS-DNA, improves their toxicity profile, but in some cases also reduces their efficacy or duration of effect. Reducing PS content improved the acute tolerability of ASOs in both mice and sheep. We show that this acute toxicity is not mediated by the major nucleic acid sensing innate immune pathways. Formulating ASOs with calcium ions before injecting into the CNS further improved their tolerability, but through a mechanism at least partially distinct from the reduction of PS content. Overall, our work identifies and quantifies an understudied aspect of oligonucleotide toxicology in the CNS, explores its mechanism, and presents platform-level medicinal chemistry approaches that improve tolerability of this class of compounds.

## INTRODUCTION

Antisense oligonucleotides (ASOs) are emerging as a promising class of therapeutics for neurological diseases.^1–4^ The groundbreaking ASO nusinersen was approved by the FDA in 2016 to treat spinal muscular atrophy.^5–12^ Nusinersen operates through a splice-switching mechanism and is fully modified with 2’-O-methoxyethyl sugars and phosphorothioate linkages. Nusinersen showed an excellent safety and efficacy profile in clinical trials. Another class of ASOs is designed to recruit RNase H to cleave target RNA and thus silence gene expression. These so-called gapmer ASOs require a stretch (“gap”) of 8-10 non-sugar-modified DNA nucleotides in the middle to enable RNase H recruitment. In contrast to nusinersen, early clinical evaluation of gapmer ASOs in the CNS faced dose-limiting toxicity resulting in failed clinical trials. Nevertheless, after these false starts, a more advanced generation of gapmer ASOs are in clinical development for neurological diseases including Huntington’s disease,^13–15^ Alzheimer’s disease,^16^ Parkinson’s disease,^17^ and amyotrophic lateral sclerosis (ALS).^4, 18–20^ These compounds show promising biomarker efficacy with reasonable toxicity profiles.

Looking at the chemical modification patterns used in early clinical trials vs RNase H-active compounds currently in the clinic, a major difference is the number of PS backbone modifications. In early trials, gapmer ASOs were fully PS modified (similar to nusinersen), while current variants of the compounds use a mixed backbone where up to six linkages are substituted back to phosphodiester. In parallel in our own work, during the development of gapmer ASOs for *C9ORF72*-driven ALS,^21^ we observed a range of transient motor phenotypes most severe within the first 1-3 hours after intracerebroventricular administration of the ASOs to mice. We were able to reduce these effects by reducing the backbone PS content. We carried out the current study to explore the generality of this phenomenon, explore its mechanism, and understand how it is affected by chemical modifications to both phosphates and sugars.

Using a quantitative scoring assay for ASO-induced motor phenotypes, we now show that the acute motor phenotypes produced by PS-modified gapmer ASOs in the brain can be minimized by using ASOs containing a combination of phosphorothioate (PS) and unmodified phosphodiester (PO) linkages. Interestingly, the ASO-induced motor phenotypes are profoundly affected by the sugar modifications used: PS-DNA ASOs were the most toxic, while ASOs composed entirely of 2’-O-substituted RNA (2’-O-methoxyethyl or 2’-O-methyl RNA) were less toxic.

We also now present experimental results that shed light on the mechanism underlying the observed motor phenotypes. We show that they are not mediated by the major nucleic acid sensing innate immune pathways, do not produce long-term toxicity and are observed in both small (mouse) and large (sheep) brains. The toxicity profile of *both* fully PS and mixed-backbone (PO/PS) ASOs can be improved by exposing ASOs to calcium-containing buffers before injection, indicating that PS-modified oligonucleotides induce a local CSF disbalance in divalent ion composition. Finally, we show that mixed (PO/PS) backbone ASOs with *in vivo* gene silencing efficacy comparable with full PS ASOs can be engineered, defining the clear steps towards development of highly active and safe ASOs for other neurodegenerative disorders.

Progress in chemical modification of oligonucleotides has been profoundly important in enabling clinical success.^22–24^ Further improvements in modification and formulation of ASOs, as well as increased mechanistic understanding of the factors defining efficacy and toxicity, is essential to expand the therapeutic use of gapmer ASOs in the CNS.

## RESULTS

### EvADINT scoring assay for acute behavioral toxicity

After administration of ASOs into the CNS, we observed dose-dependent acute behavioral toxicity that varied from lethargy, lack of responsiveness and ataxia to hyperactivity, seizures, and, in extreme cases, death. This behavioral neurotoxicity was most striking in the first 1-3 hours after administration. Even severely affected mice, unless they died, recovered fully by 24h and showed no further adverse effects. To describe our studies of acute toxicity in a robust way, we needed a way to quantify this acute toxicity.

Various protocols have been used to quantitate behavioral toxicity under the umbrella of a “Functional Observational Battery” (FOB), with variations for both acute and longitudinal neurotoxicity (reviewed in ref ^25^. Regulatory documents such as the OECD guideline for neurotoxicity testing in rodents^26^ do not provide assay details that are well aligned with the transient toxicity seen after ASO administration to mice.

We therefore developed a scoring assay optimized to quantify the transient motor phenotypes induced after intracerebral oligonucleotide injection. We call this assay EvADINT (Evaluation of Acute Drug-Induced NeuroToxicity). In this assay, we monitored mice at multiple time points over 24 hours after injection, assigning a score for various parameters as described in Table 1. If a mouse died, it was given a score of 75. Seizures, hyperactivity and other atypical motor behavior were scored depending on their severity. The rest of the score was allotted based on how much time elapsed before the animal was able to resume various aspects of normal mouse behavior. We weighted the observed phenotypes according to their apparent severity. For example, sternal posture was weighted more heavily than normal grooming since mice must be able to right themselves to carry out most other aspects of normal mouse behavior, and because maintenance of sternal posture is simpler than eating, walking or grooming. For similar intuitive reasons, seizures and death were weighted more heavily than the other factors such as latency to resume normal mouse behavior(s). We varied the relative weightings of different factors in our assay – death, seizures, and the various other behavioral observations – and observed that the relative scores of different ASOs were very similar in all cases. Thus, the EvADINT scoring assay is robust to variation in the precise weights assigned to each factor.

**Table 1.**
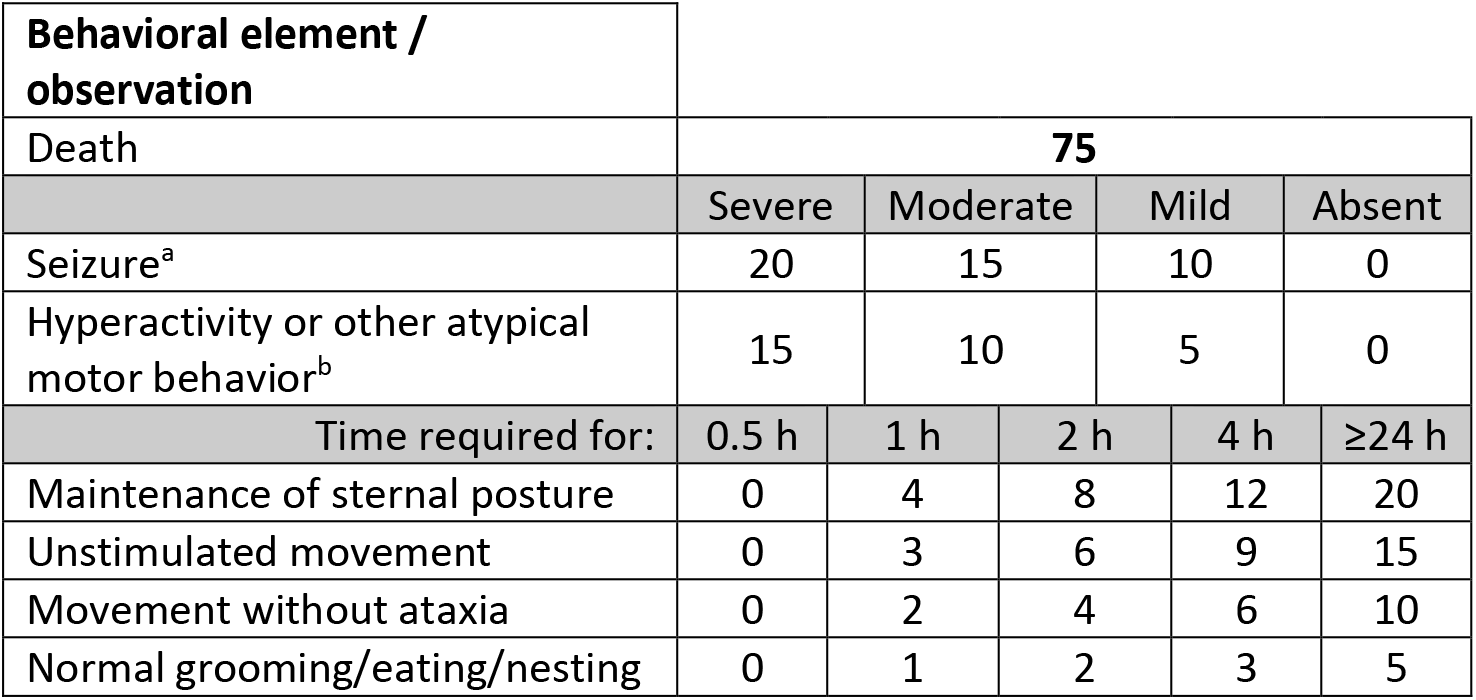
Breakdown of scoring for the EvADINT system for quantification of acute motor phenotypes. Examples of observed phenotypes (movies and corresponding scores), and all actual mouse scores for all figures are given in the supporting information. Personnel were blinded to ASO group during the scoring. ^a^Severity of seizures was ranked as follows: A *severe* seizure had a duration of >30min and/or with constant or high intensity muscle contractions. A *moderate* seizure had a 10-30 min duration and moderate intensity muscle contractions or rapid and repetitive synchronous twitching, accompanied by apparent short-term loss of consciousness. A *mild* seizure lasted <10min and featured low intensity muscle contractions or short & infrequent bursts. ^b^Severity of hyperactivity or other atypical motor behavior was ranked as follows. Severe: >30min duration, popcorning/jumping, constant. Moderate: 10-30min duration, slight hopping, other atypical motor behavior. Mild: <10min duration, uncoordinated movements, twitching.

### Reduced phosphorothioate content and formulation with Ca^2+^ reduces acute toxicity

In our parallel work on the development of ASOs that target the sense and antisense transcripts from the ALS/FTD gene *C9ORF72*, we found that reducing the number of phosphorothioate-modified linkages between pairs of 2’-*O*-methoxyethyl (MOE)-modified ribonucleotides improved tolerability without loss of efficacy or duration of effect.^21^ A similar mixed backbone design is now in clinical development.^15^ Thus our lead backbone design, and the mixed backbone design that we focus on exclusively in this study, contains a single PS linkage at each end of the ASO, followed by three PO linkages and then PS linkages throughout the remaining (central) portion of the ASO (sequences and modification patterns are shown in Table 2). With the EvADINT scoring system established, we set out to quantify this observation and explore its generality and mechanism. We first established that our previous result was reproducible under quantitative, blinded conditions. Thus, reducing the phosphorothioate content of a *C9ORF72*-targeted ASO led to increased tolerability in the CNS (Figure 1A).

**Table 2.**
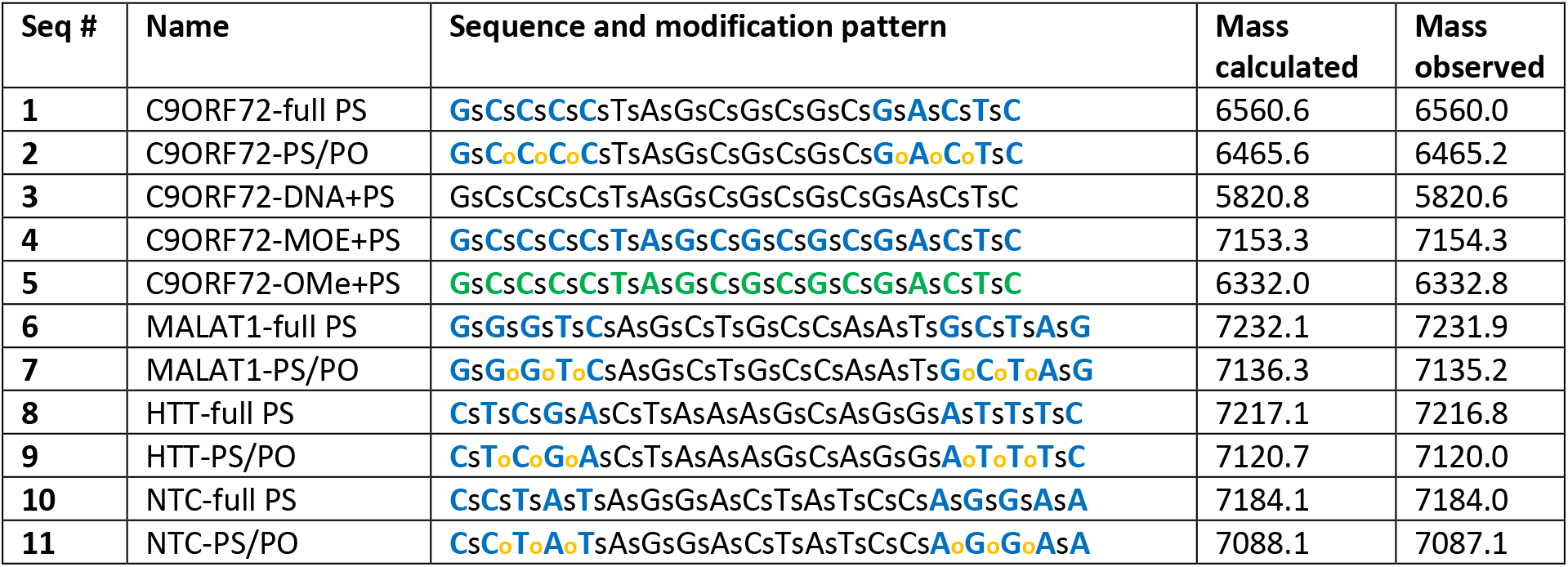
Sequences used in this study. Bold blue: MOE. Bold green: 2’-OMe. Black: DNA. Lower case letters s and o refer to phosphorothioate and phosphodiester linkages, respectively. NTC: non-target control ASO.

**Figure 1.**
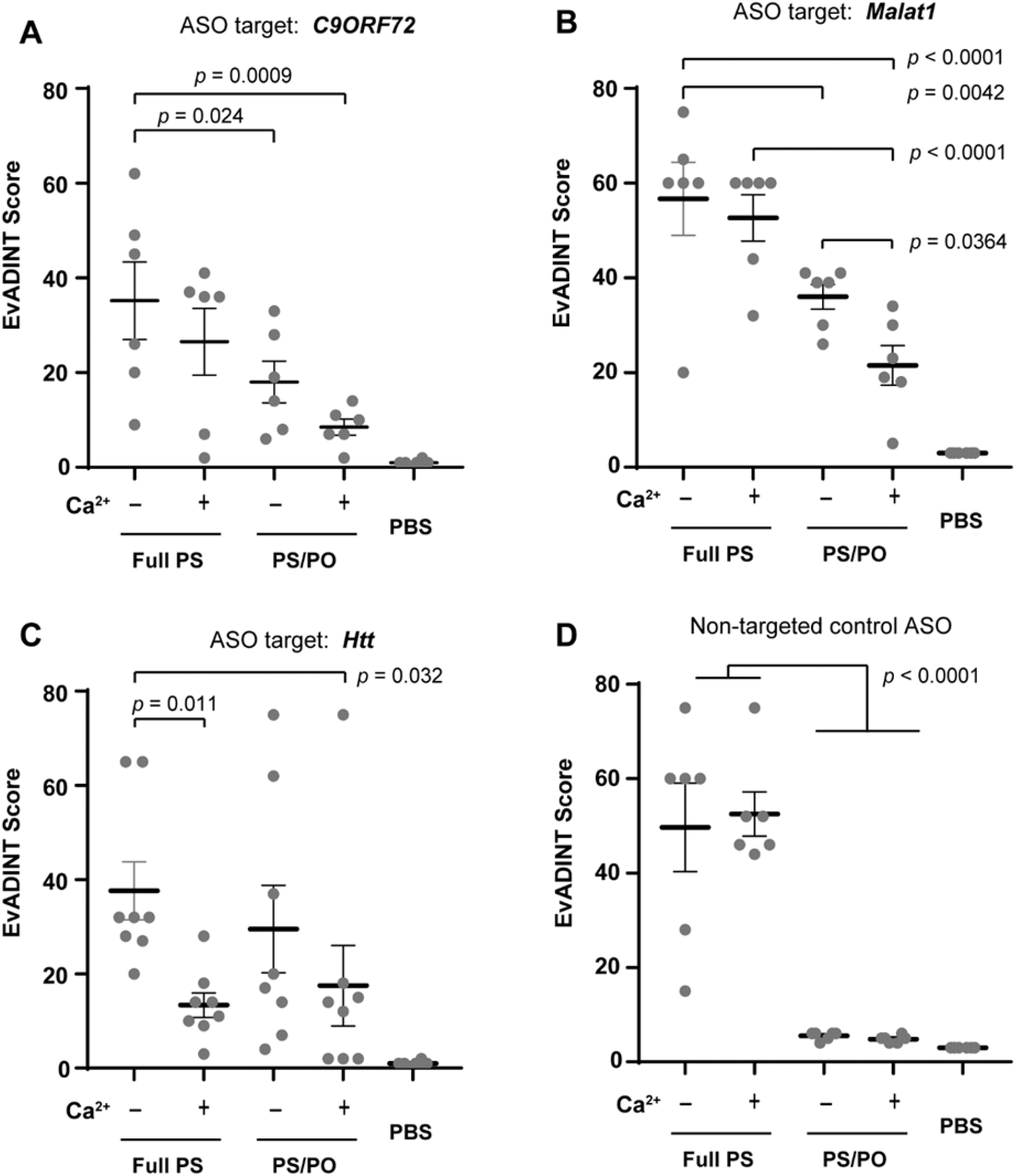
The tolerability of ASOs administered into the CNS is improved by modestly reducing the backbone PS content and by formulating with Ca^2+^ ions before injecting. Mice were injected ICV with 35 nmol of each ASO in 10 μL PBS (or with 10 μL PBS as control) and behavior was scored by a blinded investigator over the following 24h using the EvADINT rubric. Sequences targeting (A) *C9ORF72*, (B) *Malat1*, or (C) *Htt*, or (D) a non-targeting control ASO, showed improvements in acute tolerability upon reducing the PS content, formulating with Ca^2+^ ions, or both. Each data point represents the EvADINT score from one mouse; n = 6–8; error bars represent SEM. P-values are calculated using one-way ANOVA within GraphPad Prism software and represent per-comparison error rates. Full sequences and modification patterns are given in Table 2.

Phosphorothioate groups are more acidic than phosphates, and thus more anionic at physiological pH, and a greater share of the negative charge is concentrated on the sulfur atom. We wondered whether the more anionic character of phosphorothioate-modified ASOs was increasing their tendency to chelate divalent cations from CSF. During ICV injections, to keep volumes small, the ASO concentration is high (typically 1-4 mM). This is comparable to or higher than the concentration of Ca^2+^ in CSF, which is 1.3–1.4 mM^27^ – and moreover, each polyanionic ASO could potentially chelate multiple Ca^2+^ ions. We wondered whether reducing the PS content of our ASOs was reducing toxicity simply by reducing their tendency to chelate Ca^2+^ ions.

To explore whether the neurotoxicity we observed could be explained by Ca^2+^ chelation, we tested ASOs that had been pre-saturated with Ca^2+^ before injection. We chose this pre-saturation approach because we did not know exactly how much Ca^2+^ each ASO would chelate from the solution. We were concerned that if we simply suspended our ASOs in buffer containing a physiological concentration of Ca^2+^, it might be insufficient to compensate for the chelation abilities of ASOs at these high concentrations. And on the other hand, we were concerned that injecting ASOs in buffer containing higher-than-physiological concentrations of Ca^2+^ might lead to hypercalcemia-mediated toxicity. Thus, after HPLC purification of ASOs, we transferred each ASO to a 3kDa-cutoff Amicon ultrafiltration cartridge, and washed with a solution containing 20 mM Ca^2+^, twice with water, and once with PBS.^28^ We then resuspended the ASO in PBS and proceeded to injection. The intervening water washes are important to prevent the irreversible precipitation of calcium phosphate resulting from excess Ca^2+^ in the presence of phosphate buffer.

For this study, we chose a moderately high dose of 35 nmol ASO per mouse (equivalent to about 10mg/kg), somewhat higher than the dose typically required for effective gene silencing. Under these conditions, for our *C9ORF72*-targeted ASO, we observed that pre-saturation with Ca^2+^ led to a robust improvement in acute tolerability in the CNS (Figure 1A). Interestingly, the improvement in tolerability occurred for both the full PS and the mixed backbone ASOs. Thus, the best-tolerated ASO was the compound with reduced PS content and which had also been pre-saturated with Ca^2+^.

These C9ORF72 ASOs targeted the human transcript; the acute motor phenotypes were observed whether the ASOs had a target (as in the transgenic C9ORF72 mouse models we used in the parallel work on therapeutic development for C9ORF72)^21^ or not (as in the wild type mice used here).

To test whether these principles applied to other targeting and non-targeting sequences, we synthesized ASOs targeting the noncoding RNA *Malat1* and the Huntingtin (*Htt*) mRNA (Sequences **6**–**9**, Table 2). We synthesized versions of these sequences in both full PS and mixed backbone formats, and in all cases compared simple formulation in PBS with formulation in PBS after calcium pre-saturation. We injected these compounds into mice (ICV) and observed motor phenotypes using the EvADINT assay (Figure 1B-C). For the *Malat1*-targeting ASO (Figure 1B), we saw the same pattern as for the *C9ORF72*-targeted ASO – namely, there was a substantial improvement in tolerability upon reducing the PS content, and both the full PS and mixed backbone ASOs showed further improvement upon formulation with Ca^2+^.

For the *Htt*-targeted ASO, we saw higher toxicity and broader variability in tolerability across the groups (Figure 1C). The improvement in tolerability of this mixed-backbone design was less clear than for the other sequences. However, the improvement in tolerability from Ca^2+^ formulation was robust in the context of both backbone variants. Thus, there may be a sequence-dependence to optimal backbone design, and it is clear that reducing the PS content in this way does not reduce the need to select good sequences.

Finally, we synthesized non-targeting control ASOs (Sequences **10** and **11**, Table 2) and formulated them in the same way as the first three sequences. We injected these ASOs into mice and scored neurotoxicity using the EvADINT assay (Figure 1D). The patterns observed for this sequence are slightly different – in this case, calcium formulation had little or no effect on toxicity, while the reduction of PS content showed a dramatic improvement in toxicity.

Thus, each of the four sequences we tested showed a robust improvement in toxicity following a modest reduction in PS content, Ca^2+^ formulation, or both. The ASO-induced motor phenotypes were dose dependent across multiple sequences and design patterns (Supporting Figure S1). It is interesting that across all sequences, higher toxicity was often accompanied with higher inter-animal variability. That is, there appear to be important animal-to-animal variations in the actualization of this toxicity.

### Sugar 2’-modifications reduce acute motor phenotypes

Nusinersen, approved to treat spinal muscular atrophy,^9–12^ contains a fully phosphorothioate backbone. Yet it is well tolerated in the CNS^8^ and has received FDA approval.^5, 6^ This compound is modified at each nucleotide with MOE; it is not a gapmer and does not require a stretch of DNA because it functions to redirect splicing rather than recruiting RNase H. We therefore wondered whether fully-phosphorothioate-modified ASOs containing different sugar modifications might show acute motor phenotypes to a different extent.

To study this question, we synthesized fully-phosphorothioate ASOs modified *at every nucleotide* with DNA, 2’-O-methyl RNA, or MOE, respectively (in contrast to the gapmer designs used in Figure 1). We suspended these in PBS, injected them ICV at 35 nmol/mouse and scored acute motor phenotypes using the EvADINT assay. The fully DNA oligonucleotide **3** was dramatically more toxic than the two oligonucleotides containing 2’-modifications at every nucleotide (**4** and **5**; Figure 2). The two 2’-modified versions showed a dramatic reduction in motor phenotypes. We also carried out the comparison of full MOE with full DNA for a sequence targeting Malat1 and saw the same dramatic difference in toxicity (data not shown: the Malat1 experiment was carried out before we had established the quantitative EvADINT assay, but the clear difference we observed strongly suggests that the impact of sugar modification on toxicity is not sequence specific.)

**Figure 2.**
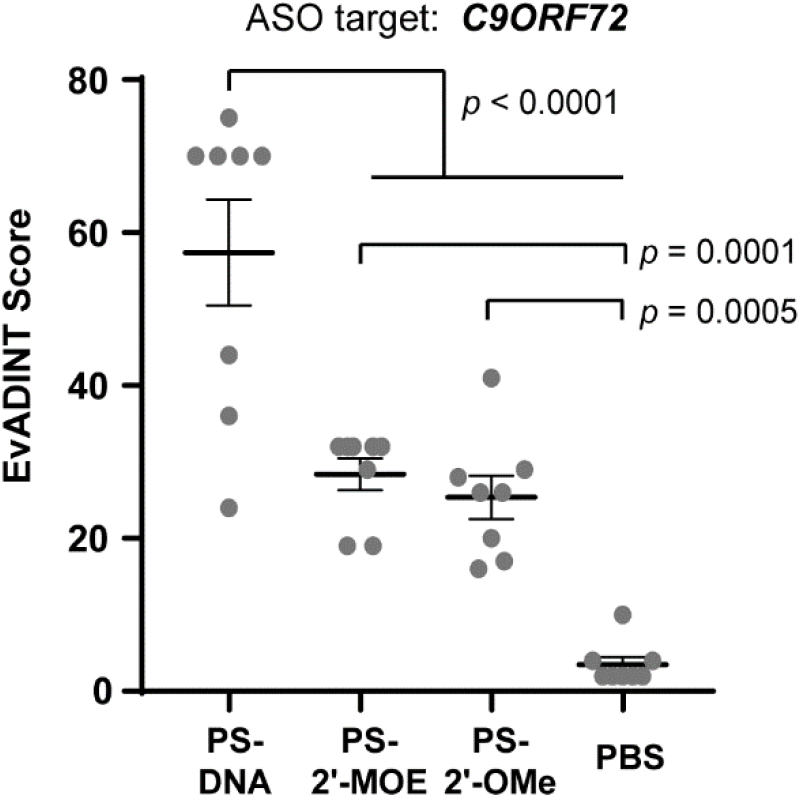
Fully PS ASOs containing 2’-modifications at each position are less toxic than those containing DNA at each position. We injected 35 nmol of each ASO in PBS to the right lateral ventricle of mice, and recorded behavioral outcomes according to the EvADINT rubric. Each data point represents the EvADINT score from one mouse; n = 8; error bars represent SEM. P-values are calculated using one-way ANOVA within GraphPad Prism software and represent per-comparison error rates. Full sequences and modification patterns are given in Table 2.

### PS-ASO-induced acute neurotoxicity is not mediated by the major nucleic acid sensing innate immune pathways

The acute toxicity we observe is most intense in the first hour after injection of mice. This timing suggested to us that the acute toxicity is not mediated by the innate immune system, since innate immune responses to nucleic acid stimuli typically do not peak until several hours after stimulation.^29, 30^

To confirm in a more direct way whether innate immune responses could play a role, we directly evaluated whether there was any contribution to this neurotoxicity from signaling through Toll-like receptors (TLRs) 3, 7, or 9, or the CGAS-STING pathway. First, we evaluated mouse behavior after injecting a 50 μg dose of poly(I:C), a compound and dose known to induce potent innate immune stimulation through TLR3 or MDA-5.^29,31^ Mice treated with poly(I:C) showed no evidence of the acute motor phenotypes seen with PS-modified ASOs, with EvADINT scores comparable to the buffer-only control mice (Figure 3A).

**Figure 3.**
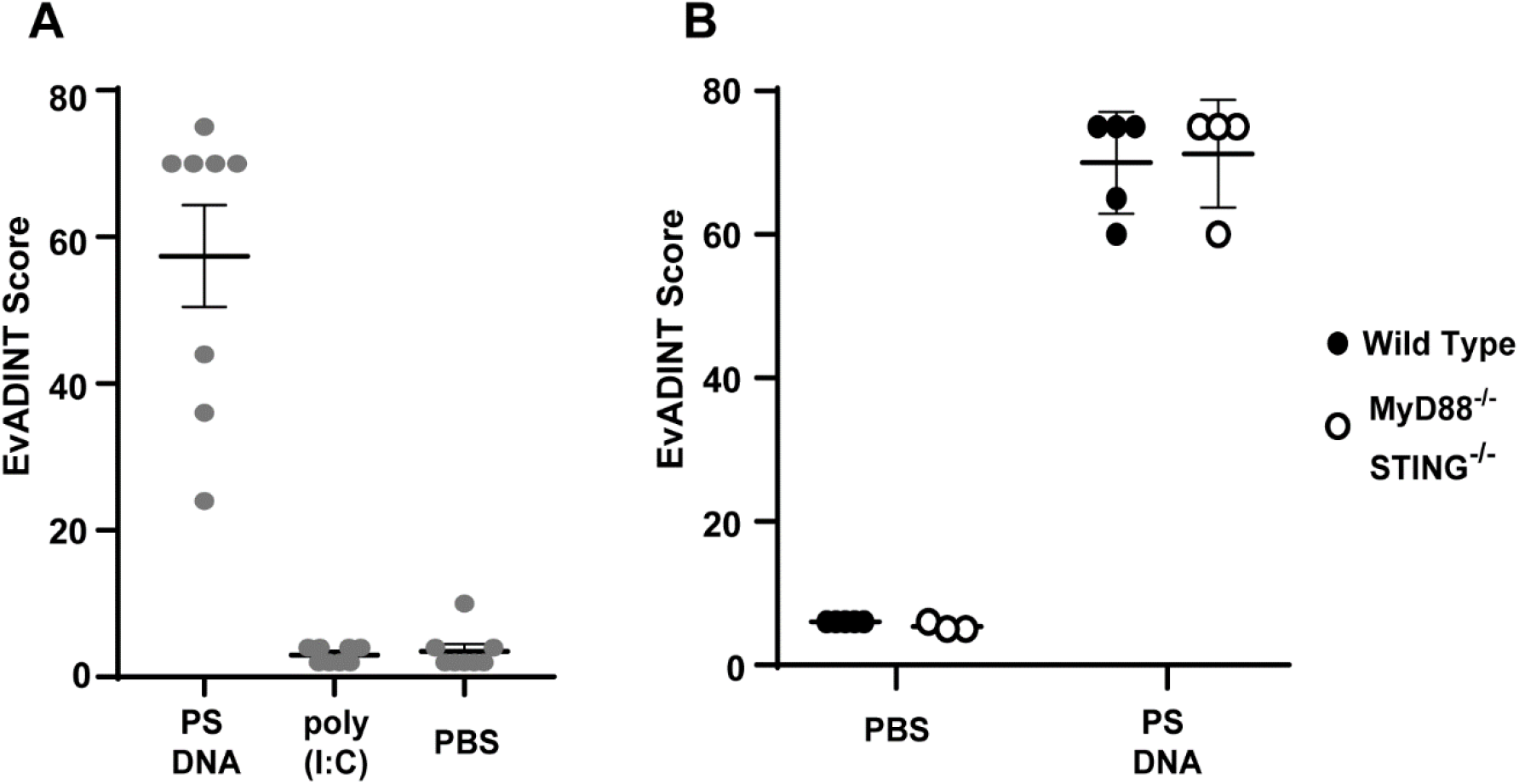
The acute neurotoxicity we observe is not mediated by toll-like receptors 3, 7, or 9, by MDA-5, or by the cGAS-STING pathway. (A) The highly immunogenic compound poly(I:C) produces none of the behavioral toxicity seen for the PS-containing ASOs after ICV injection, showing that the toxicity is not signaling through TLR3 or MDA-5. We injected 35 nmol of PS-DNA or 50 μg of poly(I:C) in PBS to the right lateral ventricle of mice, and recorded behavioral outcomes according to the EvADINT rubric. (B) Double knockout MyD88^−/−^ STING^−/−^ mice (hollow dots) show an identical response to wild type mice (filled dots), showing that the acute toxicity is not mediated by TLR7, TLR9 or the CGAS-STING pathway. In both panels, each data point represents the EvADINT score from one mouse; error bars represent SEM. All PS-DNA samples were significantly different from all other samples in both panels (One-way ANOVA, *p* < 0.0001). Full sequences and modification patterns are given in Table 2.

We also evaluated the role of other endosomal receptors TLRs 7 and 9 as well as the cytosolic DNA sensing cGAS-STING pathway by obtaining mice lacking both MyD88 and STING (Myd88^−/−^ STING^−/−^ double knockout mouse). This strain lacks signaling components for both endosomal as well as cytosolic nucleic acid sensing pathways. TLRs 7 and 9 signal through MyD88, whereas STING functions downstream of cGAS following cytosolic DNA sensing. Injection of a PS-DNA ASO into these double knockout mice showed an identical response relative to a background-matched control mouse, confirming that the toxicity is not mediated by any of these nucleic acid sensors (Figure 3B).

Taken together, these experiments provide evidence that the acute toxicity which is the focus of this work is not mediated by the major nucleic acid sensing innate immune pathways. Of course, this finding does not preclude a role for other oligonucleotide-induced innate immune responses in the brain at longer timepoints, as described by other authors.^32^

### The improved tolerability of mixed backbone ASOs applies to large brains

We wondered whether the acute toxicity we observed was an artifact of the small brain size of mice. This would make the concern significantly less relevant to researchers interested in therapeutic development of ASOs. To test whether the phenomenon applied to larger brains, we injected two sheep with fully phosphorothioate ASO (C9ORF72-full PS, Sequence **1**), and four sheep with the mixed backbone analogue of the same sequence (C9ORF72-PS/PO, Sequence **2**).

Direct intrathecal injection, the route used for patients receiving ASO therapeutics, is not practical in sheep because of difficulty accessing the intrathecal compartment and because CSF tends to be expelled from the site where the dura is punctured, leading to poor uptake of ASO. Therefore, we used a technique whereby a microcatheter was threaded up though the intrathecal space and the ASO was delivered directly into the cisterna magna (see Methods). In all cases successful microcatheter navigation was performed into cisterna magna. Both intracisternal contrast injection and cone beam computed tomography confirmed the correct catheter position prior to ASO injection. Contrast material opacification was seen in the cisterna magna, around the cerebellum and in the upper cervical spinal canal. No complication was observed in relation to catheter navigation or contrast injection.

None of the four sheep that were given the mixed backbone ASO (C9ORF72-PS/PO, Sequence **2**) showed evidence of abnormal motor phenotypes. In contrast, both sheep that were given fully phosphorothioate ASO (C9ORF72-full PS, Sequence **1**) showed hindlimb weakness and gait instability (wobbliness) within the first 24 hours. Thus, the acute toxicity of fully phosphorothioate ASOs is not specific to mice but also applies to large brains. The ASOs were given in Lactated Ringer’s solution, a calcium-containing diluent readily available at USP-grade, which confirms that the toxicity improvement mediated by reducing PS content is at least partly distinct from the question of Ca^2+^ chelation, in larger brains (in this case, sheep) as in mice as described above.

### Impact on efficacy of reducing phosphorothioate content and Ca^2+^ formulation

We previously observed for *C9ORF72*-targeted ASOs that the mixed backbone strategy did not reduce potency or efficacy as long as the phosphodiester linkages were between MOE nucleotides on both sides. In contrast, when we modified the linkage between a MOE nucleotide and a deoxynucleotide, the potency dropped dramatically.^21^ Thus, in this paper we have focused on a single mixed backbone design, in which the three internal linkages in each MOE wing were PO, and all other positions were PS. To study the effect on efficacy of this backbone design across these sequences, we compared the gene silencing of ASOs against HTT and MALAT1 in their full PS and mixed backbone versions. To be able to discriminate between compound efficacy, we chose a non-saturating dose for this element of our study: we therefore injected 15 nmol of each ASO ICV, and harvested brains after 3 weeks.

We found that both HTT and MALAT1-targeted ASOs significantly reduced their target mRNA expression, but there was a trend to reduced efficacy in mixed backbone format relative to the fully PS analogues (Figure 4A-B). It is not clear whether this results from a reduction in nuclease stability or a reduction in cellular uptake, since the PS linkage contributes to both factors. Nevertheless, this finding suggests that there is merit in exploring alternate backbone architectures, including next-generation mixed backbone designs,^33^ that might allow improvements in acute toxicity while maintaining or improving potency and efficacy.

**Figure 4.**
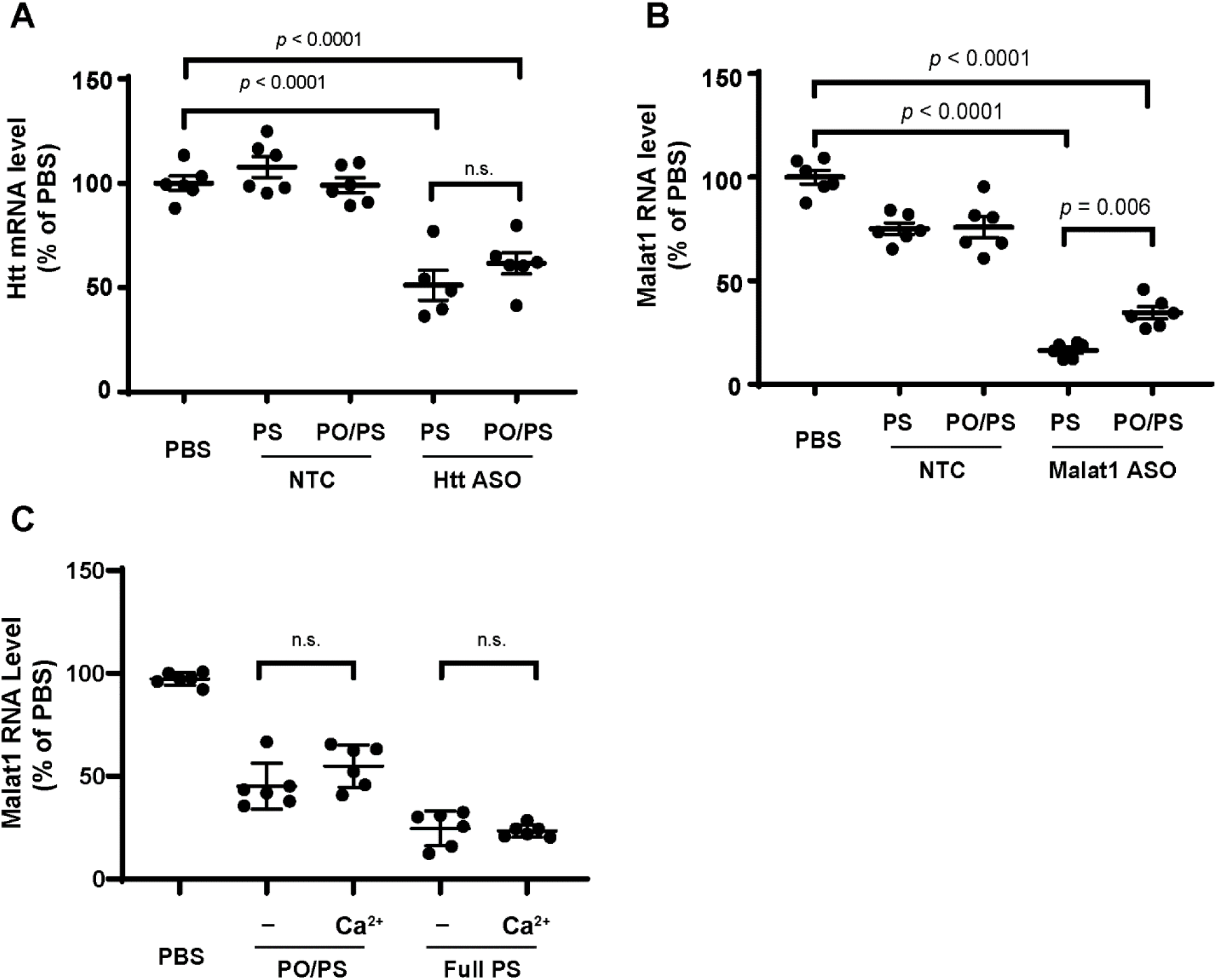
Effect of phosphorothioate reduction and calcium formulation on ASO efficacy. (A,B) Silencing of (A) *Htt* and (B) *Malat1* RNA three weeks after a 15 nmol dose of either full PS of mixed backbone (PO/PS) ASOs. Each data point represents one mouse; n=6 mice. (C) Formulation with calcium does not affect the gene silencing efficacy of ASOs. ASOs were resuspended and delivered in PBS, with or without first saturating the ASOs with calcium (see methods). In all panels, statistical significance was evaluated by one-way ANOVA followed by Tukey’s multiple comparisons test. Sequence numbers relate to the sequences and modification patterns given in Table 2.

Next, we tested whether calcium pre-saturation affected ASO efficacy. We prepared ASOs by dissolving in PBS, either with or without a Ca^2+^ pre-saturation step, and injected them ICV into mice. In the context of either a full PS or mixed PO/PS backbone, we observed that gene silencing efficacy was not affected by the presence of the Ca^2+^ pre-saturation step (Figure 4C).

## DISCUSSION

Over the past few years, an increasing number of papers have described the use of gapmer ASOs in the CNS. Many of these studies employ ASOs containing full-PS backbones^4, 13, 34–36^ or those for which the modification pattern is not clearly disclosed.^16, 18, 37, 38^ Other recent papers do describe ASOs containing a mixture of PS and PO linkages for use in the CNS.^15, 17, 19, 20, 36, 39–42^ However, to the best of our knowledge, no comparative data on the neurotoxicology of these mixed-backbone ASOs relative to fully-PS ASOs has been presented, nor has the relationship of sugar modification with acute motor phenotypes been previously known or the underlying mechanism explored before now.

The mechanism for the acute neurotoxicity of PS-modified ASOs is likely multifactorial. We initially wondered whether the more polyanionic nature of PS linkages was increasing chelation of divalent ions by ASOs. Since divalent cations, and Ca^2+^ in particular, are key for synaptic signaling and neurotransmission,^43, 44^ such depletion of divalent cations from CSF would be expected to produce acute toxicity that would be expected to last until homeostasis of Ca^2+^ concentration in CSF was restored. Ca^2+^ chelation has been responsible for unexpected toxicities in other classes of drugs.^45^ Supporting a role for this mechanism, pre-saturating ASOs with Ca^2+^ before injection improved their tolerability. Other groups have observed ASO-mediated Ca^2+^ chelation in cultured neurons.^46^ However, mixed backbone strategies improved toxicity even in the presence of such divalent ions, which suggests that Ca^2+^ chelation cannot fully explain the acute motor phentoypes induced by PS-modified gapmer ASOs.

Our experiments document that ASO-induced transient motor phenotypes are not a downstream consequence of innate immune signaling through the major nucleic-acid-sensing immune pathways. Treatment with poly(I:C) did not recapitulate the acute toxicity induced by treatment with PS-modified ASOs, suggesting that the PS-ASO-induced motor phenotypes do not result from TLR3- or MDA-5-mediated effects. And experiments in Myd88^−/−^ STING^−/−^ mice confirmed that PS-ASO-induced motor phenotypes are not mediated by TLR7, TLR9, or cGAS/STING.

At least part of the acute neurotoxicity of PS-ASOs is likely to be mediated by protein binding, for example, to cell surface receptors involved in neuronal signaling. PS-DNA shows extensive binding to a variety of proteins,^47–50^ including cell-surface and trafficking proteins.^51–54^ The origins of the high protein binding of PS-oligos were recently explored with a structural study.^55^ After systemic administration, an earlier generation of mixed-backbone modification ASOs showed reduced binding to proteins such as complement pathway members and clotting factors.^56–59^ An explanation of the acute motor phenotypes related to protein binding is also consistent with the sugar modification data presented above, since PS-MOE reduces nonspecific protein binding relative to PS-DNA;^60^ for example, recent work showed that PS-MOE bound various plasma proteins with 3 to 50-fold lower affinity relative to PS-DNA.^61^ In another recent study, PS-2’OMe gapmers of two different sequences showed about 2.5-fold lower affinity protein binding than PS-DNA of the same sequence.^55^

The acute motor phenotypes discussed in this work do not appear to be related to other types of ASO-induce toxicity – such as liver toxicity or immune stimulation. For example, in ongoing work in our group, we have come across sequences that are well-tolerated in terms of acute motor phenotypes, but still exhibit liver toxicity, and vice versa. The timing of these effects is also very different: with acute motor phenotypes strongest in the first hour (perhaps driven by binding to cell-surface receptors as discussed above), innate immune stimulation peaking at 1-2 days (driven by binding to toll-like receptors and cytosolic nucleic acid sensors), and liver toxicity evident from 1 day to several days after dosing (driven by factors including mislocalization of paraspeckle proteins to nucleoli^62^).

This paper has focused on a single mixed-backbone design, and the impact of replacing three PS linkages within each MOE wing with their corresponding (unmodified) PO linkages. It is striking that a relatively small reduction in PS content makes such a dramatic difference in acute toxicity for multiple sequences. Our work on *C9ORF72* showed that this design not only maintained but improved potency relative to the full PS version.^21^ Other investigators have also disclosed that some ASOs show increased potency when a subset of PS linkages are replaced by PO linkages.^63^ Nevertheless, this design sometimes leads to a modest loss in potency, as shown in Figure 4.

The well-tolerated nature of <u>fully 2’-modified</u> ASOs in the CNS (Figure 2) has allowed the rapid development and FDA approval of nusinersen^5–9^ as well as the first personalized ASO drug, milasen.^64^ However, recruitment of RNase H requires the presence either of DNA or a DNA analogue, typically as a gapmer design.^65, 66^ The fact that PS-DNA shows higher acute toxicity than the corresponding 2’-modified nucleotides (Figure 2) suggests that a major area of focus for nucleic acid chemists should be the development of sugar or phosphate-modified DNA analogues that elicit robust RNase H cleavage while reducing the incidence of motor phenotypes when administered to the CNS. The development of next-generation mixed-backbone ASOs is an active area of research in our group.^33, 67^ In the meantime, researchers can implement mixed backbone designs such as those described here, along with calcium formulation, to improve the therapeutic index of gapmer ASOs for clinical use in the CNS.

## METHODS

### Oligonucleotides

All oligonucleotides were synthesized using ABI 394 or Akta OligoPilot synthesizers using standard methods. Phosphoramidites were purchased from ChemGenes and diluted to 0.1 M in acetonitrile. Sulfurization was accomplished using DDTT (0.1M, ChemGenes). Benzylthiotetrazole (0.25 M in acetonitrile, TEDIA) was used as activator. All cytosine residues were 5-methylcytosine.

Oligonucleotides were deprotected by treatment with concentrated aqueous ammonia at 55°C for 16h then concentrated and purified by ion-exchange HPLC (eluting with 30% acetonitrile in water containing increasing gradients of NaClO_4_) or ion-pairing reverse-phase HPLC (eluting with aqueous triethylammonium acetate containing increasing gradients of acetonitrile). All oligonucleotides were characterized by LCMS.

After HPLC purification, we carried out final desalting and buffer equilibration using ultrafiltration (Amicon centrifugal filters, 3-kDa molecular weight cutoff, Millipore). For oligonucleotides administered in PBS, we placed the oligonucleotide in the Amicon filter and washed with 2 changes of PBS. To saturate calcium-binding sites within oligonucleotides, we placed the purified oligonucleotides in the Amicon filter, washed them twice with a 20 mM solution of CaCl_2_, twice with water, and once with PBS, before resuspending the ASOs in PBS (Note, in follow up work after the completion of this study, we found that a lower concentration of CaCl_2_ was preferable to minimize compound loss during this wash step, which is an issue for certain sequences). For the oligonucleotides used for sheep studies, we did not carry out calcium saturation in this manner but rather resuspended in USP-grade Lactated Ringers Solution (LRS, which contains 1.3 mM Ca^2+^).

### ICV administration of oligonucleotides in mice

Mouse studies were carried out under UMass Medical School IACUC protocol A-2551. FVB/NCI mice (7-13 weeks old) were anaesthetized by intraperitoneal injection of fentanyl/midazolam/dexmedetomidine (0.1, 5, and 0.25 mg/kg, respectively, as a solution in sterile saline). Anaesthetized animals were transferred to a Kopf small animal stereotaxic frame, ear bars placed and a hand-warmer was placed underneath the animal to preserve core body-temperature. Ophthalmic lubricant was placed over each eye and the fur covering the skull was removed. The scalp was aseptically prepared by thorough alternate swabbing with betadine and 70% isopropanol (3x each) and allowed to dry before a medial incision was made to expose the skull. The periosteum was dried with a sterile cotton swab and the syringe containing oligonucleotide was moved to bregma. A point was marked on the skull 1 mm dextrolateral and 0.4 mm posterior from bregma and a 0.6–0.8-mm diameter hole drilled at this location. The tip of the needle was advanced 2 mm ventrally through this hole into the lateral ventricle and after a 2 minute wait, the oligonucleotide was injected over a period of 25 seconds (10 μL total injection volume). The needle was left in place for 3 minutes post-injection, then removed and the skin closed with 5-0 vicryl suture. An intraperitoneal injection of fluemazenil/atipamezole (0.5 mg/kg and 5 mg/kg respectively, in sterile saline) was used to reverse injected anesthetic agents. Buprenorphine was also injected for analgesia (0.3 mg/kg, SC). Animals were removed from the stereotaxic frame and allowed to recover in a warm cage, food and gel were provided, and the animals were observed periodically over the next 24 hours according to the rubric laid out in Table 1.

For ICV injections in the Myd88^−/−^ STING^−/−^ mice (C57BL/6 background) and corresponding C57BL/6 controls, we used the protocol described above with slightly adjusted coordinates for the ICV injection (1mm dextrolateral, 2mm posterior, 3mm ventral.)

### STING/MyD88 double knockout mice

Myd88^−/−^ mice on C57BL/6 background^68^ were obtained from S. Akira (Osaka University, Osaka, Japan). STING^−/−^ mice on C57BL/6 background^69^ were originally from G. Barber (University of Miami, Florida) and obtained from D. Stetson (University of Washington, Seattle). The two strains were intercrossed to generate Myd88^−/−^ STING^−/−^ double knockouts. The mice were bred and maintained under pathogen–free conditions in our animal facility. The Myd88 and STING deficiencies were confirmed by performing PCR on DNA obtained after digesting a tail snip. For Myd88, specific primer pairs were used to distinguish the WT or knockout allele in two separate reactions. Reaction 1 with primer sequences AGC CTC TAC ACC CTT CTC TTC TCC ACA and AGA CAG GCT GAG TGC AAA CTT GTG CTG was used to detect the WT band at 1000 bp and reaction 2 with primer pairs AGC CTC TAC ACC CTT CTC TTC TCC ACA and ATC GCC TTC TAT CGC CTT CTT GAC GAG were used to detect KO band at 1000 bp. For STING, reaction 1 with primer sequences AGA ACG GAC AGC CAG TAA GTA TAC AG and CAA TGC TCT CAT AGC CTT CAC TAT C was used to detect the WT band at 375 bp and reaction 2 with primer pairs AAC TTC CTG ACT AGG GGA GGA GTA G and CAA TGC TCT CAT AGC CTT CAC TAT C was used to detect the KO band at 470 base pairs.

### ASO administration in sheep

Sheep studies were carried out under UMass Medical School IACUC protocol A-2593. Jacob sheep were fasted overnight in preparation for surgery. A 20G catheter was placed and secured in the jugular vein, blood (5 mL) was drawn from the catheter for analysis, and the catheter was flushed with saline (0.9% NaCl). We administered buprenorphine (0.01 mg/kg IM), acepromazine (0.05 mg/kg IM) and glycopyrrolate (0.01 mg/kg IM) 30 min prior to induction of anesthesia. An intravenous cocktail of ketamine (6 mg/kg) and diazepam (0.3 mg/kg) was administered to induce anesthesia, followed by ketoprofen (2.2. mg/kg SQ) as analgesic and cefazolin (22 mg/kg IV) to minimizes any risk of infection. The animal was intubated, and a stomach tube was placed to prevent rumen gas pressure build up. Anesthesia was maintained using vaporized isoflurane (1.5–3.5% in oxygen). Sheep were positioned in lateral recumbency in an Allura Xper FD20 X-ray system (Philips Medical Systems, Best, Netherlands). A 19-gauge Tuohy needle was inserted in the lumbosacral (L7-S1) intrathecal space and then ~5 mL of CSF was removed. Using the Touhy needle as entry point, a straight tip microcatheter (Excelsior SL-10; Stryker Neurovascular) was inserted through the lumen of the needle to access the intrathecal space. The microcatheter was navigated into the cisterna magna with an assistance of a 0.014” wire under fluoroscopic guidance. The wire had a slight curve on the tip (Synchro Guidewire, Stryker Neurovascular) to avoid any nerve or vascular structure damage. A microcatheter contrast injection (1 mL of Omnipaque 240 mgI/ml) was injected and the pattern of contrast material distribution was visualized prior to the injection of ASO solution (2 mg/kg in ~3 mL of Lactated Ringers Solution; for comparison, the mouse doses of 15-35 nmol/mouse equate to about 4-10 mg/kg). Cone beam computed tomography (Allura Xper FD20 X-ray system) imaging was performed to confirm the final microcatheter position in the cisterna magna in relation to the nerve and vascular structures. At the end of the procedure the Touhy needle was withdrawn and the microcatheter was removed.

### EvADINT scoring system for acute neurotoxicity

After ICV administration of ASOs to mice, a blinded investigator ranked the behavior of mice at multiple timepoints using the rubric laid out in Table 1. If a mouse died within the first 24 hours, its score was assigned to be 75; otherwise it was the sum of all other scores. Seizures, if observed, were scored based on severity; hyperactive or spastic behavior was also scored based on severity and included twitching, uncontrolled movement such as “popcorning” and other atypical motor phenotypes. Besides these elements, the score was based on the time elapsed until mice resumed normal posture and behavior; for example, if a mouse required more than 1 hour but less than 2 hours to be able to right itself (resume and maintain sternal posture) it would be given a score of 8. Each mouse was individually scored. Examples of scoring, with corresponding videos, are provided in Supporting Table S1. The breakdown of scoring for each mouse is provided in Supporting Tables S2-S7.

### Evaluation of gene silencing in the CNS

#### For comparison of backbones on gene silencing efficacy

Mice were euthanized at 3 weeks post-treatment by cervical dislocation and the brain was immediately removed into ice-cold PBS. The brain was placed in a brain matrix (Braintree scientific) and the most rostral 3 mm discarded. A 1-mm slice was then taken and each side homogenized independently. The tissue was suspended in Affymetrix homogenizing solution containing proteinase K and mechanically dissociated using a Quiagen Tissuelyser with a 2mm tungsten carbide bead. The tubes were then incubated in a water bath at 65 °C until all tubes appeared transparent. Tubes were centrifuged (16,000 xG, 15 minutes) and supernatant transferred to a 96 well plate for storage at −80. *Htt, Malat1* and *Ppib* RNA levels were quantified using the QuantiGene 2.0 assay kit (Affymetrix, QS0011) as previously described.^70^

#### For studying the effect calcium formulation on gene silencing efficacy

Mice were euthanized at 2 weeks post-injection via IP administration of 0.1mL of 390 mg/mL pentobarbital sodium and the brain and spinal cord were immediately removed into ice-cold PBS. A 2mm section of the lumbar spinal cord was cut and placed in an Eppendorf tube in −80°C. The brain was placed in a brain matrix and the most rostral 3mm discarded. A 2-mm slice was taken, and the cortical section removed and placed in an Eppendorf tube in −80°C. Tissue was homogenized in TRI-reagent using a Qiagen TissueLyser and 1 μg of RNA was reverse transcribed using High-Capacity cDNA Reverse Transcription kit (Life Technologies) per the manufacturer’s protocol. qRT-PCR was carried out using iTaq Supermix (Bio-Rad) on Bio-Rad CFX-96 real time machine using gene-specific primers: *Malat1* Primer1: 5’ CTC CAA CAA CCA CTA CTC CAA 3’; Primer2: 5’ GTA CTG TTC CAA TCT GCT GCT A 3’; probe: /56-FAM/TCA TAC TCC /ZEN/AGT CGC GTC ACA ATG C/3IABkFQ/. For *Ppib* (internal control), Primer1: 5’CCG TAG TGC TTC AGC TTG A 3’; Primer2: 5’ AGC AAG TTC CAT CGT GTC ATC 3’; Probe: /56-FAM/TGC TCT TTC /ZEN/CTC CTG TGC CAT CTC /3IABkFQ/.

## SUPPORTING INFORMATION / DATA AVAILABILITY

Supporting information is available in the online version of this file, and includes the following: Supporting Figure S1 shows the dose responsiveness of acute motor phenotypes. Supporting Table S1 and associated movies show examples of mouse phenotypes and corresponding assigned scores. Tables S2-S10 provide a breakdown of the EvADINT scoring for each mouse and each treatment.

## ACKNOWLEDGEMENTS

We are grateful to Art Levin for suggesting that we explore the idea of Ca^2+^ formulation as a means of reducing acute neurotoxicity. We thank Nina Bishop, Andrew Coles, Neil Aronin, Miguel Esteves and Matthew Gounis for help with the sheep study. We thank Anastasia Khvorova for critical feedback on the manuscript. This work was funded by the Ono Pharmaceutical Foundation (Breakthrough Science Award to JKW), the Friedreich’s Ataxia Research Alliance (Grant to JKW) and the NIH (R01 NS111990 to RHB and JKW). RHB acknowledges funding from The Angel Fund for ALS Research, ALSOne, ALS Finding a Cure, the Cellucci Fund for ALS Research and the Max Rosenfeld Fund.

## AUTHOR CONTRIBUTIONS

MPM and PMK synthesized and purified oligonucleotides; MPM, JMR-B, FW and AW carried out mouse experiments; FW, JMR-B and JKW developed the EvADINT scoring assay; MPM, JMR-B, M.Marosfoi, RMK and HG-E carried out the sheep study; M.Motwani and KAF developed the knockout mice, JKW supervised the study and wrote the manuscript; all authors edited the manuscript.

## REFERENCES

1. Seguin, R., Antisense Oligonucleotides for Treatment of Neurological Diseases. In Oligonucleotide-Based Drugs and Therapeutics: Preclinical and Clinical Considerations for Development. (eds. N. Ferrari & R. Seguin) 389–409 (Wiley, Hoboken; 2018).

2. Evers, M.M., Toonen, L.J. & van Roon-Mom, W.M. (2015). Antisense oligonucleotides in therapy for neurodegenerative disorders. Adv Drug Deliv Rev 87, 90–103.

3. Wahlestedt, C. (1994). Antisense oligonucleotide strategies in neuropharmacology. Trends Pharmacol. Sci. 15, 42–46.

4. Smith, R.A., Miller, T.M., Yamanaka, K., Monia, B.P., Condon, T.P., Hung, G., Lobsiger, C.S., Ward, C.M., McAlonis-Downes, M., Wei, H. et al. (2006). Antisense oligonucleotide therapy for neurodegenerative disease. J. Clin. Invest. 116, 2290–2296.

5. Aartsma-Rus, A. (2017). FDA Approval of Nusinersen for Spinal Muscular Atrophy Makes 2016 the Year of Splice Modulating Oligonucleotides. Nucleic Acid Ther 27, 67–69.

6. Corey, D.R. (2017). Nusinersen, an antisense oligonucleotide drug for spinal muscular atrophy. Nat. Neurosci. 20, 497–499.

7. Hua, Y., Sahashi, K., Hung, G., Rigo, F., Passini, M.A., Bennett, C.F. & Krainer, A.R. (2010). Antisense correction of SMN2 splicing in the CNS rescues necrosis in a type III SMA mouse model. Genes Dev. 24, 1634–1644.

8. Passini, M.A., Bu, J., Richards, A.M., Kinnecom, C., Sardi, S.P., Stanek, L.M., Hua, Y., Rigo, F., Matson, J., Hung, G. et al. (2011). Antisense Oligonucleotides Delivered to the Mouse CNS Ameliorate Symptoms of Severe Spinal Muscular Atrophy. Sci. Trans. Med. 3, 72ra18–72ra18.

9. Rigo, F., Hua, Y., Krainer, A.R. & Bennett, C.F. (2012). Antisense-based therapy for the treatment of spinal muscular atrophy. J. Cell Biol. 199, 21–25.

10. Singh, N.N., Howell, M.D., Androphy, E.J. & Singh, R.N. (2017). How the discovery of ISS-N1 led to the first medical therapy for spinal muscular atrophy. Gene Ther. 24, 520–526.

11. Finkel, R.S., Mercuri, E., Darras, B.T., Connolly, A.M., Kuntz, N.L., Kirschner, J., Chiriboga, C.A., Saito, K., Servais, L., Tizzano, E. et al. (2017). Nusinersen versus Sham Control in Infantile-Onset Spinal Muscular Atrophy. N. Engl. J. Med. 377, 1723–1732.

12. Mercuri, E., Darras, B.T., Chiriboga, C.A., Day, J.W., Campbell, C., Connolly, A.M., Iannaccone, S.T., Kirschner, J., Kuntz, N.L., Saito, K. et al. (2018). Nusinersen versus Sham Control in Later-Onset Spinal Muscular Atrophy. N. Engl. J. Med. 378, 625–635.

13. Kordasiewicz, H.B., Stanek, L.M., Wancewicz, E.V., Mazur, C., McAlonis, M.M., Pytel, K.A., Artates, J.W., Weiss, A., Cheng, S.H., Shihabuddin, L.S. et al. (2012). Sustained therapeutic reversal of Huntington’s disease by transient repression of huntingtin synthesis. Neuron 74, 1031–1044.

14. Tabrizi, S., Leavitt, B., Kordasiewicz, H., Czech, C., Swayze, E., Norris, D.A., Baumann, T., Gerlach, I., Schobel, S., Smith, A. et al. (2018). Effects of IONIS-HTTRx in Patients with Early Huntington’s Disease, Results of the First HTT-Lowering Drug Trial (CT.002). Neurology 90.

15. Tabrizi, S.J., Leavitt, B.R., Landwehrmeyer, G.B., Wild, E.J., Saft, C., Barker, R.A., Blair, N.F., Craufurd, D., Priller, J., Rickards, H. et al. (2019). Targeting Huntingtin Expression in Patients with Huntington’s Disease. N. Engl. J. Med. 380, 2307–2316.

16. DeVos, S.L., Miller, R.L., Schoch, K.M., Holmes, B.B., Kebodeaux, C.S., Wegener, A.J., Chen, G., Shen, T., Tran, H., Nichols, B. et al. (2017). Tau reduction prevents neuronal loss and reverses pathological tau deposition and seeding in mice with tauopathy. Sci. Trans. Med. 9, eaag0481.

17. Zhao, H.T., John, N., Delic, V., Ikeda-Lee, K., Kim, A., Weihofen, A., Swayze, E.E., Kordasiewicz, H.B., West, A.B. & Volpicelli-Daley, L.A. (2017). LRRK2 Antisense Oligonucleotides Ameliorate α-Synuclein Inclusion Formation in a Parkinson’s Disease Mouse Model. Mol. Ther. Nucl. Acids 8, 508–519.

18. Jiang, J., Zhu, Q., Gendron, Tania F., Saberi, S., McAlonis-Downes, M., Seelman, A., Stauffer, Jennifer E., Jafar-nejad, P., Drenner, K., Schulte, D. et al. (2016). Gain of Toxicity from ALS/FTD-Linked Repeat Expansions in *C9ORF72* Is Alleviated by Antisense Oligonucleotides Targeting GGGGCC-Containing RNAs. Neuron 90, 535–550.

19. Becker, L.A., Huang, B., Bieri, G., Ma, R., Knowles, D.A., Jafar-Nejad, P., Messing, J., Kim, H.J., Soriano, A., Auburger, G. et al. (2017). Therapeutic reduction of ataxin-2 extends lifespan and reduces pathology in TDP-43 mice. Nature 544, 367.

20. McCampbell, A., Cole, T., Wegener, A.J., Tomassy, G.S., Setnicka, A., Farley, B.J., Schoch, K.M., Hoye, M.L., Shabsovich, M., Sun, L. et al. (2018). Antisense oligonucleotides extend survival and reverse decrement in muscle response in ALS models. J. Clin. Invest. 128, 3558–3567.

21. Tran, H., Moazami, M.P., Yang, H., McKenna-Yasek, D., Douthwright, C., Pinto, C., Metterville, J., Shin, M., Sanil, N., Dooley, C. et al. (2021). Potent mixed backbone antisense oligonucleotide safely suppresses expression of mutant c9orf72 transcripts and polydipeptides: First in human pilot study. Research Square (preprint) DOI: 10.21203/rs.3.rs-211236/v1.

22. Khvorova, A. & Watts, J.K. (2017). The chemical evolution of oligonucleotide therapies of clinical utility. Nature Biotechnology 35, 238–248.

23. Watts, J.K., The medicinal chemistry of antisense oligonucleotides. In Oligonucleotide-Based Drugs and Therapeutics. (eds. N. Ferrari & R. Seguin) 39–90 (Wiley, Hoboken; 2018).

24. Wan, W.B. & Seth, P.P. (2016). The medicinal chemistry of therapeutic oligonucleotides. J. Med. Chem. 59, 9645–9667.

25. Moser, V.C. (2011). Functional Assays for Neurotoxicity Testing. Toxicol. Pathol. 39, 36–45.

26. OECD GUIDELINE FOR THE TESTING OF CHEMICALS (1997). Test No. 424: Neurotoxicity Study in Rodents DOI 10.1787/20745788.

27. Murphy, V.A., Smith, Q.R. & Rapoport, S.I. (1986). Homeostasis of brain and cerebrospinal fluid calcium concentrations during chronic hypo- and hypercalcemia. J. Neurochem. 47, 1735–1741.

28. In more recent unpublished work, we have identified that 5 mM calcium is a better concentration for this saturation protocol, since for some sequences 20 mM calcium leads to reduced solubility.

29. Town, T., Jeng, D., Alexopoulou, L., Tan, J. & Flavell, R.A. (2006). Microglia Recognize Double-Stranded RNA via TLR3. The Journal of Immunology 176, 3804–3812.

30. Jiang, M., Zhang, S., Yang, Z., Lin, H., Zhu, J., Liu, L., Wang, W., Liu, S., Liu, W., Ma, Y. et al. (2018). Self-Recognition of an Inducible Host lncRNA by RIG-I Feedback Restricts Innate Immune Response. Cell 173, 906–919.e913.

31. Kato, H., Takeuchi, O., Sato, S., Yoneyama, M., Yamamoto, M., Matsui, K., Uematsu, S., Jung, A., Kawai, T., Ishii, K.J. et al. (2006). Differential roles of MDA5 and RIG-I helicases in the recognition of RNA viruses. Nature 441, 101–105.

32. Toonen, L.J.A., Casaca-Carreira, J., Pellise-Tintore, M., Mei, H., Temel, Y., Jahanshahi, A. & van Roon-Mom, W.M.C. (2018). Intracerebroventricular Administration of a 2’-O-Methyl Phosphorothioate Antisense Oligonucleotide Results in Activation of the Innate Immune System in Mouse Brain. Nucleic Acid Ther 28, 63–73.

33. Debacker, A.J., Sharma, V.K., Meda Krishnamurthy, P., O’Reilly, D., Greenhill, R. & Watts, J.K. (2019). Next-Generation Peptide Nucleic Acid Chimeras Exhibit High Affinity and Potent Gene Silencing. Biochemistry 58, 582–589.

34. Ostergaard, M.E., Southwell, A.L., Kordasiewicz, H., Watt, A.T., Skotte, N.H., Doty, C.N., Vaid, K., Villanueva, E.B., Swayze, E.E., Bennett, C.F. et al. (2013). Rational design of antisense oligonucleotides targeting single nucleotide polymorphisms for potent and allele selective suppression of mutant Huntingtin in the CNS. Nucleic Acids Res 41, 9634–9650.

35. DeVos, S.L., Goncharoff, D.K., Chen, G., Kebodeaux, C.S., Yamada, K., Stewart, F.R., Schuler, D.R., Maloney, S.E., Wozniak, D.F., Rigo, F. et al. (2013). Antisense Reduction of Tau in Adult Mice Protects against Seizures. J. Neurosci. 33, 12887–12897.

36. Meng, L., Ward, A.J., Chun, S., Bennett, C.F., Beaudet, A.L. & Rigo, F. (2014). Towards a therapy for Angelman syndrome by targeting a long non-coding RNA. Nature 518, 409.

37. Friedrich, J., Kordasiewicz, H.B., O’Callaghan, B., Handler, H.P., Wagener, C., Duvick, L., Swayze, E.E., Rainwater, O., Hofstra, B., Benneyworth, M. et al. (2018). Antisense oligonucleotide-mediated ataxin-1 reduction prolongs survival in SCA1 mice and reveals disease-associated transcriptome profiles. JCI insight 3, e123193.

38. Luo, X., Fitzsimmons, B., Mohan, A., Zhang, L., Terrando, N., Kordasiewicz, H. & Ji, R.-R. (2018). Intrathecal administration of antisense oligonucleotide against p38α but not p38β MAP kinase isoform reduces neuropathic and postoperative pain and TLR4-induced pain in male mice. Brain. Behav. Immun. 72, 34–44.

39. Sztainberg, Y., Chen, H.M., Swann, J.W., Hao, S., Tang, B., Wu, Z., Tang, J., Wan, Y.W., Liu, Z., Rigo, F. & Zoghbi, H.Y. (2015). Reversal of phenotypes in MECP2 duplication mice using genetic rescue or antisense oligonucleotides. Nature 528, 123–126.

40. Moore, L.R., Rajpal, G., Dillingham, I.T., Qutob, M., Blumenstein, K.G., Gattis, D., Hung, G., Kordasiewicz, H.B., Paulson, H.L. & McLoughlin, H.S. (2017). Evaluation of Antisense Oligonucleotides Targeting ATXN3 in SCA3 Mouse Models. Mol. Ther. Nucl. Acids 7, 200–210.

41. Mohan, A., Fitzsimmons, B., Zhao, H.T., Jiang, Y., Mazur, C., Swayze, E.E. & Kordasiewicz, H.B. (2017). Antisense oligonucleotides selectively suppress target RNA in nociceptive neurons of the pain system and can ameliorate mechanical pain. Pain 159, 139–149.

42. Ling, K.K., Jackson, M., Alkam, D., Liu, D., Allaire, N., Sun, C., Kiaei, M., McCampbell, A. & Rigo, F. (2018). Antisense-mediated reduction of EphA4 in the adult CNS does not improve the function of mice with amyotrophic lateral sclerosis. Neurobiol. Dis. 114, 174–183.

43. Berridge, M.J. (1998). Neuronal Calcium Signaling. Neuron 21, 13–26.

44. Brini, M., Cali, T., Ottolini, D. & Carafoli, E. (2014). Neuronal calcium signaling: function and dysfunction. Cell. Mol. Life Sci. 71, 2787–2814.

45. Bosakowski, T. & Levin, A.A. (1987). Comparative acute toxicity of chlorocitrate and fluorocitrate in dogs. Toxicology and Applied Pharmacology 89, 97–104.

46. Olson, R.E., Cacace, A.M., Hagedorn, P., Hog, A.M., Jensen, M.L., Nielsen, N.F., Li, D., Brown, J.M. & Mercer, S.E. (2016). Tau antisense oligomers and uses thereof. Patent application WO 2016/126995 A1.

47. Brown, D.A., Kang, S.H., Gryaznov, S.M., DeDionisio, L., Heidenreich, O., Sullivan, S., Xu, X. & Nerenberg, M.I. (1994). Effect of phosphorothioate modification of oligodeoxynucleotides on specific protein binding. Journal of Biological Chemistry 269, 26801–26805.

48. Lebedeva, I. & Stein, C.A. (2001). Antisense oligonucleotides: promise and reality. Annual Review of Pharmacology and Toxicology 41, 403–419.

49. Levin, A.A., Yu, R.Z. & Geary, R.S., Basic Principles of the Pharmacokinetics of Antisense Oligonucleotide Drugs. In Antisense Drug Technology: Principles, Strategies, and Applications, Edn. 2nd. (ed. S.T. Crooke) 183–215 (CRC Press, Boca Raton; 2008).

50. Geary, R.S. (2009). Antisense oligonucleotide pharmacokinetics and metabolism. Expert Opin Drug Met 5, 381–391.

51. Weidner, D.A., Valdez, B.C., Henning, D., Greenberg, S. & Busch, H. (1995). Phosphorothioate oligonucleotides bind in a non sequence-specific manner to the nucleolar protein C23/nucleolin. FEBS Letters 366, 146–150.

52. Stein, C.A., Wu, S., Voskresenskiy, A.M., Zhou, J.F., Shin, J., Miller, P., Souleimanian, N. & Benimetskaya, L. (2009). G3139, an anti-Bcl-2 antisense oligomer that binds heparin-binding growth factors and collagen I, alters in vitro endothelial cell growth and tubular morphogenesis. Clin. Cancer Res. 15, 2797–2807.

53. Miller, C.M., Donner, A.J., Blank, E.E., Egger, A.W., Kellar, B.M., Østergaard, M.E., Seth, P.P. & Harris, E.N. (2016). Stabilin-1 and Stabilin-2 are specific receptors for the cellular internalization of phosphorothioate-modified antisense oligonucleotides (ASOs) in the liver. Nucleic Acids Research 44, 2782–2794.

54. Wang, S., Allen, N., Vickers, T.A., Revenko, A.S., Sun, H., Liang, X.H. & Crooke, S.T. (2018). Cellular uptake mediated by epidermal growth factor receptor facilitates the intracellular activity of phosphorothioate-modified antisense oligonucleotides. Nucleic Acids Res 46, 3579–3594.

55. Hyjek-Skladanowska, M., Vickers, T.A., Napiorkowska, A., Anderson, B.A., Tanowitz, M., Crooke, S.T., Liang, X.H., Seth, P.P. & Nowotny, M. (2020). Origins of the Increased Affinity of Phosphorothioate-Modified Therapeutic Nucleic Acids for Proteins. J. Am. Chem. Soc. 142, 7456–7468.

56. Agrawal, S., Rustagi, P.K. & Shaw, D.R. (1995). Novel enzymatic and immunological responses to oligonucleotides. Toxicology Letters 82-83, 431–434.

57. Zhang, R., Iyer, R.P., Yu, D., Tan, W., Zhang, X., Lu, Z., Zhao, H. & Agrawal, S. (1996). Pharmacokinetics and tissue disposition of a chimeric oligodeoxynucleoside phosphorothioate in rats after intravenous administration. J. Pharmacol. Exp. Ther. 278, 971–979.

58. Zhao, Q., Temsamani, J., Iadarola, P.L., Jiang, Z. & Agrawal, S. (1996). Effect of different chemically modified oligodeoxynucleotides on immune stimulation. Biochem. Pharmacol. 51, 173–182.

59. Agrawal, S., Jiang, Z., Zhao, Q., Shaw, D., Sun, D. & Saxinger, C. (1997). Mixed-Backbone Oligonucleotides Containing Phosphorothioate and Methylphosphonate Linkages as Second Generation Antisense Oligonucleotide. Nucleosides Nucleotides 16, 927–936.

60. Henry, S.P., Kim, T.-W., Kramer-Stickland, K., Zanardi, T.A., Fey, R.A. & Levin, A.A., Toxicologic Properties of 2’-O-Methoxyethyl Chimeric Antisense Inhibitors in Animals and Man. In Antisense Drug Technology: Principles, Strategies and Applications. (ed. S.T. Crooke) 327–363 (CRC Press, Boca Raton; 2007).

61. Gaus, H.J., Gupta, R., Chappell, A.E., Ostergaard, M.E., Swayze, E.E. & Seth, P.P. (2019). Characterization of the interactions of chemically-modified therapeutic nucleic acids with plasma proteins using a fluorescence polarization assay. Nucleic Acids Res 47, 1110–1122.

62. Shen, W., De Hoyos, C.L., Migawa, M.T., Vickers, T.A., Sun, H., Low, A., Bell, T.A., 3rd, Rahdar, M., Mukhopadhyay, S., Hart, C.E. et al. (2019). Chemical modification of PS-ASO therapeutics reduces cellular protein-binding and improves the therapeutic index. Nat. Biotechnol. 37, 640–650.

63. Ferrari, N. & Seguin, R. (2012). Oligonucleotide inhibitors with chimeric backbone and 2-amino-2’-deoxyadenosine. PCT application PCT/CA2012/000169.

64. Kim, J., Hu, C., Moufawad El Achkar, C., Black, L.E., Douville, J., Larson, A., Pendergast, M.K., Goldkind, S.F., Lee, E.A., Kuniholm, A. et al. (2019). Patient-Customized Oligonucleotide Therapy for a Rare Genetic Disease. N. Engl. J. Med. 381, 1644–1652.

65. Damha, M.J., Wilds, C.J., Noronha, A., Brukner, I., Borkow, G., Arion, D. & Parniak, M.A. (1998). Hybrids of RNA and arabinonucleic acids (ANA and 2’F-ANA) are substrates of Ribonuclease H. J. Am. Chem. Soc. 120, 12976–12977.

66. Verbeure, B., Lescrinier, E., Wang, J. & Herdewijn, P. (2001). RNase H mediated cleavage of RNA by cyclohexene nucleic acid (CeNA). Nucleic Acids Research 29, 4941–4947.

67. Sharma, V.K., Singh, S.K., Krishnamurthy, P.M., Alterman, J.F., Haraszti, R.A., Khvorova, A., Prasad, A.K. & Watts, J.K. (2017). Synthesis and biological properties of triazole-linked locked nucleic acid. Chem. Commun. 53, 8906–8909.

68. Adachi, O., Kawai, T., Takeda, K., Matsumoto, M., Tsutsui, H., Sakagami, M., Nakanishi, K. & Akira, S. (1998). Targeted disruption of the MyD88 gene results in loss of IL-1- and IL-18-mediated function. Immunity 9, 143–150.

69. Ishikawa, H. & Barber, G.N. (2008). STING is an endoplasmic reticulum adaptor that facilitates innate immune signalling. Nature 455, 674–678.

70. Coles, A.H., Osborn, M.F., Alterman, J.F., Turanov, A.A., Godinho, B.M., Kennington, L., Chase, K., Aronin, N. & Khvorova, A. (2016). A High-Throughput Method for Direct Detection of Therapeutic Oligonucleotide-Induced Gene Silencing In Vivo. Nucleic Acid Ther 26, 86–92.

