## Supporting Information for "Quantifying and Mitigating Motor Phenotypes Induced by Antisense Oligonucleotides in the Central Nervous System"

**Contents:**

**Figure S1.** Dose-responsiveness of acute motor phenotypes over two sequences and modification patterns.

**Table S1.** Descriptions of movies of mice exhibiting abnormal motor behavior (the corresponding movie files are given as separate supporting files.)

**Tables S2-S10.** Raw EvADINT data from each mouse, arranged in order of the main text figures.

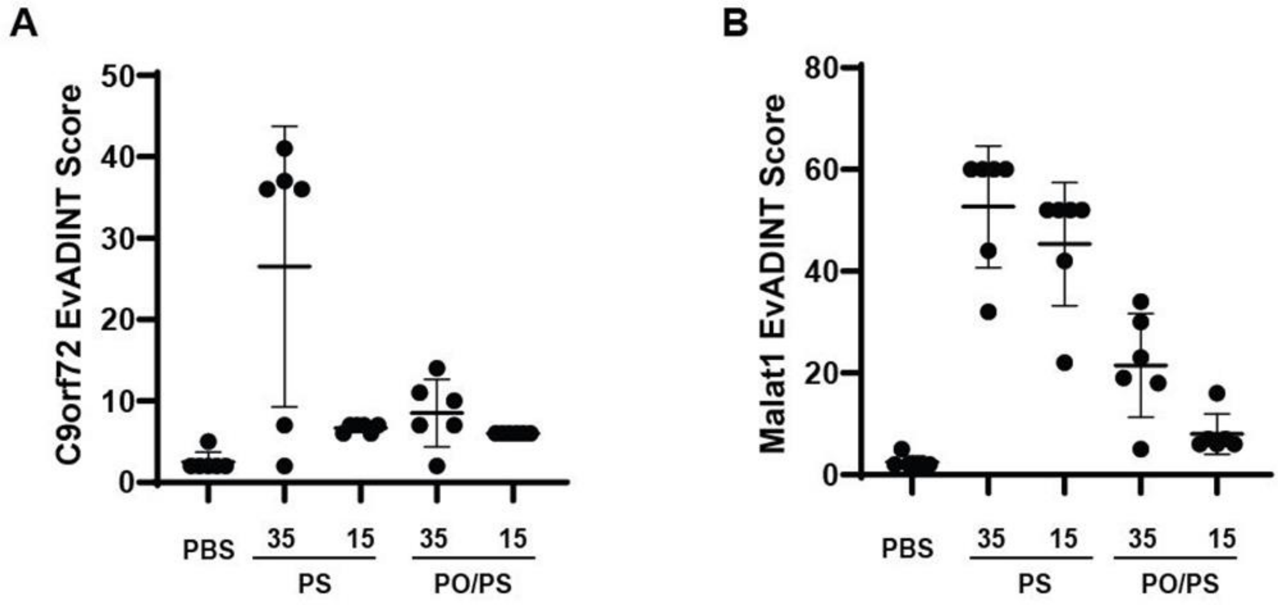


**Figure S1.** The acute neurotoxicity of ASOs is dose-dependent. Mice were injected ICV with 35 or 15 nmol of ASOs targeting (A) C9orf72 or (B) Malat1 in 10 μL PBS (or with 10 μL PBS as control) and behavior was scored by a blinded investigator over the following 24h using the EvADINT rubric. PS: fully phosphorothioate backbone, PO/PS: mixed backbone. Each data point represents the EvADINT score from one mouse; n = 6; error bars represent SEM.

**Table S1.** Descriptions of movies of mice exhibiting neurotoxic behavior (the corresponding movie files are given as separate supporting files.)

| Video | Description | Score component derived from this observation |
|---|---|---|
| Video 1: Twitching | Animal presents with a tremor / twitching / slight uncontrollable movements and with mild hyperactivity. Ranked as "other atypical motor behavior" in the EvADINT rubric, and scored as mild based on severity as well as duration ( <10 min). | 5 |
| Video 2: Ataxia | Animal presents with moderate ataxia; score in this case depends on how long it takes until the animal is able to recover normal movement. For instance, in this case the mouse was still showing ataxia at 2 h but was ambulating normally by 4 h. | 6 |
| Video 3: Seizure | Animal presents with unilateral upper and lower limb seizure-like activity, including rapid and repetitive synchronous twitching, accompanied by apparent short-term loss of consciousness. Scored as a mild seizure in the EvADINT rubric since the duration was <10 min. | 10 |
| Video 4: Atypical motor behavior | Animal presents with sustained, rapid, and repetitive unilateral head-scratching not typical of normal mouse behavior, duration <10min, scored as mild "atypical motor behavior" based on severity and duration. | 5 |
| Video 5: Hyperactivity | Animal jumps up and down alongside the cage in an uncharacteristic manner. Scored as "hyperactivity or other atypical motor behavior" and a moderate score was given based on severity and duration. | 10 |
| Video 6: Seizure | Animal presents with upper and lower limb seizure-like activity, featuring tense limbs and tremors, and accompanied by apparent loss of consciousness. Ranked as a severe seizure in the EvADINT rubric since duration was >30min. | 20 |

**The rest of the supporting information consists of tables of detailed animal-by-animal outcomes from EvADINT scoring assay experiments.** Note in the tables below (colored boxes indicating the treatment group) that “W2” is our shorthand description for the precise placement of PO and PS linkages in these mixed backbone sequences; all sequences marked “W2” are mixed PO/PS backbone.

**Table S2. Raw data from the EvADINT scoring assay for each mouse in Figure 1A (ASO targeting *C9ORF72*).**

| ASO | | | | | | Full PS | | | | | |
|---|---|---|---|---|---|---|---|---|---|---|---|
| Mouse ID | | | | | | ***41*** | ***42*** | ***43*** | ***44*** | ***45*** | ***46*** |
| Death | | | 75 | | | 0 | 0 | 0 | 0 | 0 | 0 |
| | Severe | | Moderate | | Mild | | | | | | |
| Tonic seizure | 20 | | 15 | | 10 | 0 | 20 | 15 | 0 | 0 | 15 |
| Hyperactivity or spasticity | 15 | | 10 | | 5 | 5 | 0 | 0 | 10 | 0 | 0 |
| Time required for: | 0.5 h | 1 h | 2 h | 4 h | ≥24 h | | | | | | |
| Maintenance of sternal posture | 0 | 4 | 8 | 12 | 20 | 0 | 12 | 8 | 0 | 0 | 12 |
| Unstimulated Movement | 0 | 3 | 6 | 9 | 15 | 6 | 15 | 9 | 9 | 3 | 9 |
| Movement without Ataxia | 0 | 2 | 4 | 6 | 10 | 6 | 10 | 10 | 4 | 4 | 10 |
| Normal Grooming/Eating/Nesting | 0 | 1 | 2 | 3 | 5 | 3 | 5 | 3 | 3 | 2 | 3 |
| Total Score | | | | | | **20** | **62** | **45** | **26** | **9** | **49** |

| ASO | | | | | | Full PS w/Ca2+ | | | | | |
|---|---|---|---|---|---|---|---|---|---|---|---|
| Mouse ID | | | | | | ***47*** | ***48*** | ***49*** | ***50*** | ***51*** | ***52*** |
| Death | | | 75 | | | 0 | 0 | 0 | 0 | 0 | 0 |
| | Severe | | Moderate | | Mild | | | | | | |
| Tonic seizure | 20 | | 15 | | 10 | 0 | 0 | 0 | 0 | 0 | 0 |
| Hyperactivity or spasticity | 15 | | 10 | | 5 | 10 | 0 | 5 | 5 | 0 | 0 |
| Time required for: | 0.5 h | 1 h | 2 h | 4 h | ≥24 h | | | | | | |
| Maintenance of sternal posture | 0 | 4 | 8 | 12 | 20 | 8 | 0 | 12 | 8 | 0 | 12 |
| Unstimulated Movement | 0 | 3 | 6 | 9 | 15 | 9 | 0 | 9 | 9 | 0 | 9 |
| Movement without Ataxia | 0 | 2 | 4 | 6 | 10 | 6 | 4 | 10 | 10 | 2 | 10 |
| Normal Grooming/Eating/Nesting | 0 | 1 | 2 | 3 | 5 | 3 | 3 | 5 | 5 | 0 | 5 |
| Total Score | | | | | | **36** | **7** | **41** | **37** | **2** | **36** |

| **ASO** | | | | | | **W2** | | | | | |
|---|---|---|---|---|---|---|---|---|---|---|---|
| **Mouse ID** | | | | | | ***53*** | ***54*** | ***55*** | ***56*** | ***57*** | ***58*** |
| **Death** | | | 75 | | | 0 | 0 | 0 | 0 | 0 | 0 |
| | **Severe** | | **Moderate** | | **Mild** | | | | | | |
| **Tonic seizure** | 20 | | 15 | | 10 | 0 | 0 | 0 | 0 | 0 | 0 |
| **Hyperactivity or spasticity** | 15 | | 10 | | 5 | 10 | 0 | 0 | 5 | 5 | 0 |
| **Time required for:** | **0.5 h** | **1 h** | **2 h** | **4 h** | **≥24 h** | | | | | | |
| **Maintenance of sternal posture** | 0 | 4 | 8 | 12 | 20 | 0 | 0 | 0 | 0 | 8 | 0 |
| **Unstimulated Movement** | 0 | 3 | 6 | 9 | 15 | 9 | 0 | 6 | 6 | 9 | 3 |
| **Movement without Ataxia** | 0 | 2 | 4 | 6 | 10 | 4 | 4 | 6 | 6 | 6 | 4 |
| **Normal Grooming/Eating/Nesting** | 0 | 1 | 2 | 3 | 5 | 5 | 2 | 2 | 2 | 5 | 1 |
| **Total Score** | | | | | | **28** | **6** | **14** | **19** | **33** | **8** |

| **ASO** | | | | | | **W2 w/Ca2+** | | | | | |
|---|---|---|---|---|---|---|---|---|---|---|---|
| **Mouse ID** | | | | | | ***59*** | ***60*** | ***61*** | ***62*** | ***63*** | ***64*** |
| **Death** | | | 75 | | | 0 | 0 | 0 | 0 | 0 | 0 |
| | **Severe** | | **Moderate** | | **Mild** | | | | | | |
| **Tonic seizure** | 20 | | 15 | | 10 | 0 | 0 | 0 | 0 | 0 | 0 |
| **Hyperactivity or spasticity** | 15 | | 10 | | 5 | 0 | 0 | 0 | 0 | 0 | 0 |
| **Time required for:** | **0.5 h** | **1 h** | **2 h** | **4 h** | **≥24 h** | | | | | | |
| **Maintenance of sternal posture** | 0 | 4 | 8 | 12 | 20 | 0 | 0 | 0 | 0 | 0 | 0 |
| **Unstimulated Movement** | 0 | 3 | 6 | 9 | 15 | 0 | 3 | 3 | 6 | 0 | 0 |
| **Movement without Ataxia** | 0 | 2 | 4 | 6 | 10 | 0 | 6 | 6 | 6 | 6 | 6 |
| **Normal Grooming/Eating/Nesting** | 0 | 1 | 2 | 3 | 5 | 2 | 1 | 2 | 2 | 1 | 1 |
| **Total Score** | | | | | | **2** | **10** | **11** | **14** | **7** | **7** |

| **ASO** | | | | | | **PBS** | | | | | | | |
|---|---|---|---|---|---|---|---|---|---|---|---|---|---|
| **Mouse ID** | | | | | | ***33*** | ***34*** | ***35*** | ***36*** | ***37*** | ***38*** | ***39*** | ***40*** |
| **Death** | | | 75 | | | 0 | 0 | 0 | 0 | 0 | 0 | 0 | 0 |
| | **Severe** | | **Moderate** | | **Mild** | | | | | | | | |
| **Tonic seizure** | 20 | | 15 | | 10 | 0 | 0 | 0 | 0 | 0 | 0 | 0 | 0 |
| **Hyperactivity or spasticity** | 15 | | 10 | | 5 | 0 | 0 | 0 | 0 | 0 | 0 | 0 | 0 |
| **Time required for:** | **0.5 h** | **1 h** | **2 h** | **4 h** | **≥24 h** | | | | | | | | |
| **Maintenance of sternal posture** | 0 | 4 | 8 | 12 | 20 | 0 | 0 | 0 | 0 | 0 | 0 | 0 | 0 |
| **Unstimulated Movement** | 0 | 3 | 6 | 9 | 15 | 0 | 0 | 0 | 0 | 0 | 0 | 0 | 0 |
| **Movement without Ataxia** | 0 | 2 | 4 | 6 | 10 | 0 | 0 | 0 | 0 | 0 | 0 | 0 | 2 |
| **Normal Grooming/Eating/Nesting** | 0 | 1 | 2 | 3 | 5 | 1 | 1 | 0 | 1 | 1 | 1 | 1 | 0 |
| **Total Score** | | | | | | **1** | **1** | **0** | **1** | **1** | **1** | **1** | **2** |

**Table S3. Raw data from the EvADINT scoring assay for each mouse in Figure 1B (ASO targeting *Malat1*).**

| | | | | | | | | | | | |
|---|---|---|---|---|---|---|---|---|---|---|---|
| **ASO** | | | | | | **Full PS** | | | | | |
| **Mouse ID** | | | | | | ***105*** | ***106*** | ***107*** | ***108*** | ***109*** | ***110*** |
| **Death** | 75 | | | | | | | | 75 | | |
| | **Severe** | | **Moderate** | | **Mild** | | | | | | |
| **Tonic seizure** | 20 | | 15 | | 10 | 15 | 0 | 0 | | 10 | 10 |
| **Hyperactivity or spasticity** | 15 | | 10 | | 5 | 0 | 10 | 10 | | 0 | 0 |
| **Time required for:** | **0.5 h** | **1 h** | **2 h** | **4 h** | **≥24 h** | | | | | | |
| **Maintenance of sternal posture** | 0 | 4 | 8 | 12 | 20 | 20 | 0 | 20 | | 20 | 20 |
| **Unstimulated Movement** | 0 | 3 | 6 | 9 | 15 | 15 | 3 | 15 | | 15 | 15 |
| **Movement without Ataxia** | 0 | 2 | 4 | 6 | 10 | 10 | 6 | 10 | | 10 | 10 |
| **Normal Grooming/Eating/Nesting** | 0 | 1 | 2 | 3 | 5 | 5 | 1 | 5 | | 5 | 5 |
| **Total Score** | | | | | | **65** | **20** | **60** | **75** | **60** | **60** |

| | | | | | | | | | | | |
|---|---|---|---|---|---|---|---|---|---|---|---|
| **ASO** | | | | | | **Full PS w/Ca2+** | | | | | |
| **Mouse ID** | | | | | | ***111*** | ***112*** | ***113*** | ***114*** | ***115*** | ***116*** |
| **Death** | 75 | | | | | | | | | | |
| | **Severe** | | **Moderate** | | **Mild** | | | | | | |
| **Tonic seizure** | 20 | | 15 | | 10 | 10 | 10 | 10 | 10 | 10 | 10 |
| **Hyperactivity or spasticity** | 15 | | 10 | | 5 | 0 | 0 | 0 | 0 | 0 | 0 |
| **Time required for:** | **0.5 h** | **1 h** | **2 h** | **4 h** | **≥24 h** | | | | | | |
| **Maintenance of sternal posture** | 0 | 4 | 8 | 12 | 20 | 12 | 20 | 0 | 20 | 20 | 20 |
| **Unstimulated Movement** | 0 | 3 | 6 | 9 | 15 | 9 | 15 | 9 | 15 | 15 | 15 |
| **Movement without Ataxia** | 0 | 2 | 4 | 6 | 10 | 10 | 10 | 10 | 10 | 10 | 10 |
| **Normal Grooming/Eating/Nesting** | 0 | 1 | 2 | 3 | 5 | 3 | 5 | 3 | 5 | 5 | 5 |
| **Total Score** | | | | | | **44** | **60** | **32** | **60** | **60** | **60** |

| ASO | | | | | | W2 | | | | | |
|---|---|---|---|---|---|---|---|---|---|---|---|
| Mouse ID | | | | | | *117* | *118* | *119* | *120* | *121* | *122* |
| Death | 75 | | | | | | | | | | |
| | Severe | | Moderate | | Mild | | | | | | |
| Tonic seizure | 20 | | 15 | | 10 | 0 | 0 | 0 | 0 | 0 | 0 |
| Hyperactivity or spasticity | 15 | | 10 | | 5 | 5 | 5 | 5 | 5 | 0 | 0 |
| Time required for: | 0.5 h | 1 h | 2 h | 4 h | ≥24 h | | | | | | |
| Maintenance of sternal posture | 0 | 4 | 8 | 12 | 20 | 12 | 12 | 12 | 12 | 8 | 8 |
| Unstimulated Movement | 0 | 3 | 6 | 9 | 15 | 9 | 9 | 9 | 9 | 6 | 9 |
| Movement without Ataxia | 0 | 2 | 4 | 6 | 10 | 10 | 10 | 10 | 10 | 10 | 10 |
| Normal Grooming/Eating/Nesting | 0 | 1 | 2 | 3 | 5 | 3 | 5 | 3 | 5 | 2 | 3 |
| Total Score | | | | | | **39** | **41** | **39** | **41** | **26** | **30** |

| ASO | | | | | | W2 w/Ca2+ | | | | | |
|---|---|---|---|---|---|---|---|---|---|---|---|
| Mouse ID | | | | | | *123* | *124* | *125* | *126* | *127* | *128* |
| Death | 75 | | | | | | | | | | |
| | Severe | | Moderate | | Mild | | | | | | |
| Tonic seizure | 20 | | 15 | | 10 | 0 | 0 | 0 | 0 | 0 | 0 |
| Hyperactivity or spasticity | 15 | | 10 | | 5 | 5 | 0 | 0 | 0 | 0 | 0 |
| Time required for: | 0.5 h | 1 h | 2 h | 4 h | ≥24 h | | | | | | |
| Maintenance of sternal posture | 0 | 4 | 8 | 12 | 20 | 0 | 0 | 8 | 0 | 0 | 12 |
| Unstimulated Movement | 0 | 3 | 6 | 9 | 15 | 9 | 6 | 9 | 6 | 0 | 9 |
| Movement without Ataxia | 0 | 2 | 4 | 6 | 10 | 6 | 10 | 10 | 10 | 4 | 10 |
| Normal Grooming/Eating/Nesting | 0 | 1 | 2 | 3 | 5 | 3 | 2 | 3 | 3 | 1 | 3 |
| Total Score | | | | | | **23** | **18** | **30** | **19** | **5** | **34** |

| ASO | | | | | | PBS | | | | | |
|---|---|---|---|---|---|---|---|---|---|---|---|
| Mouse ID | | | | | | *129* | *130* | *131* | *132* | *133* | *134* |
| Death | 75 | | | | | | | | | | |
| | Severe | | Moderate | | Mild | | | | | | |
| Tonic seizure | 20 | | 15 | | 10 | 0 | 0 | 0 | 0 | 0 | 0 |
| Hyperactivity or spasticity | 15 | | 10 | | 5 | 0 | 0 | 0 | 0 | 0 | 0 |
| Time required for: | 0.5 h | 1 h | 2 h | 4 h | ≥24 h | | | | | | |
| Maintenance of sternal posture | 0 | 4 | 8 | 12 | 20 | 0 | 0 | 0 | 0 | 0 | 0 |
| Unstimulated Movement | 0 | 3 | 6 | 9 | 15 | 0 | 0 | 0 | 0 | 0 | 0 |
| Movement without Ataxia | 0 | 2 | 4 | 6 | 10 | 2 | 2 | 2 | 2 | 2 | 2 |
| Normal Grooming/Eating/Nesting | 0 | 1 | 2 | 3 | 5 | 1 | 1 | 1 | 1 | 1 | 1 |
| Total Score | | | | | | **3** | **3** | **3** | **3** | **3** | **3** |

**Table S4. Raw data from the EvADINT scoring assay for each mouse in Figure 1C (ASO targeting *Htt*).**

| ASO | | | | | | Full PS | | | | | | | |
|---|---|---|---|---|---|---|---|---|---|---|---|---|---|
| Mouse ID | | | | | | ***1*** | ***2*** | ***3*** | ***4*** | ***5*** | ***6*** | ***7*** | ***8*** |
| Death | 75 | | | | | | | | | | | | |
| | Severe | | Moderate | | Mild | | | | | | | | |
| Tonic seizure | 20 | | 15 | | 10 | | | | | 15 | | | 15 |
| Hyperactivity or spasticity | 15 | | 10 | | 5 | | 10 | 10 | 10 | | | 10 | |
| Time required for: | 0.5 h | 1 h | 2 h | 4 h | ≥24 h | | | | | | | | |
| Maintenance of sternal posture | 0 | 4 | 8 | 12 | 20 | 0 | 0 | 0 | 0 | 20 | 12 | 4 | 20 |
| Unstimulated Movement | 0 | 3 | 6 | 9 | 15 | 9 | 9 | 9 | 9 | 15 | 9 | 6 | 15 |
| Movement without Ataxia | 0 | 2 | 4 | 6 | 10 | 6 | 10 | 10 | 10 | 10 | 3 | 6 | 10 |
| Normal Grooming/Eating/Nesting | 0 | 1 | 2 | 3 | 5 | 5 | 3 | 3 | 3 | 5 | 3 | 2 | 5 |
| Total Score | | | | | | **20** | **32** | **32** | **32** | **65** | **27** | **28** | **65** |

| ASO | | | | | | Full PS w/Ca2+ | | | | | | | |
|---|---|---|---|---|---|---|---|---|---|---|---|---|---|
| Mouse ID | | | | | | ***9*** | ***10*** | ***11*** | ***12*** | ***13*** | ***14*** | ***15*** | ***16*** |
| Death | 75 | | | | | | | | | | | | |
| | Severe | | Moderate | | Mild | | | | | | | | |
| Tonic seizure | 20 | | 15 | | 10 | | | | | | | | |
| Hyperactivity or spasticity | 15 | | 10 | | 5 | | | | | | | | |
| Time required for: | 0.5 h | 1 h | 2 h | 4 h | ≥24 h | | | | | | | | |
| Maintenance of sternal posture | 0 | 4 | 8 | 12 | 20 | 0 | 4 | 0 | 0 | 0 | 0 | 0 | 0 |
| Unstimulated Movement | 0 | 3 | 6 | 9 | 15 | 0 | 9 | 0 | 3 | 6 | 3 | 6 | 3 |
| Movement without Ataxia | 0 | 2 | 4 | 6 | 10 | 0 | 10 | 10 | 10 | 10 | 6 | 2 | 4 |
| Normal Grooming/Eating/Nesting | 0 | 1 | 2 | 3 | 5 | 3 | 5 | 0 | 1 | 2 | 2 | 6 | 2 |
| Total Score | | | | | | **3** | **28** | **10** | **14** | **18** | **11** | **14** | **9** |

| ASO | | | | | | W2 | | | | | | | |
|---|---|---|---|---|---|---|---|---|---|---|---|---|---|
| Mouse ID | | | | | | ***17*** | ***18*** | ***19*** | ***20*** | ***21*** | ***22*** | ***23*** | ***24*** |
| Death | 75 | | | | | | 75 | | | | | | |
| | Severe | | Moderate | | Mild | | | | | | | | |
| Tonic seizure | 20 | | 15 | | 10 | | | | | | 20 | | |
| Hyperactivity or spasticity | 15 | | 10 | | 5 | | | | | | | 15 | 5 |
| Time required for: | 0.5 h | 1 h | 2 h | 4 h | ≥24 h | | | | | | | | |
| Maintenance of sternal posture | 0 | 4 | 8 | 12 | 20 | 0 | | 0 | 0 | 0 | 12 | 4 | 0 |
| Unstimulated Movement | 0 | 3 | 6 | 9 | 15 | 6 | | 0 | 9 | 0 | 15 | 9 | 3 |
| Movement without Ataxia | 0 | 2 | 4 | 6 | 10 | 6 | | 4 | 10 | 4 | 10 | 6 | 4 |
| Normal Grooming/Eating/Nesting | 0 | 1 | 2 | 3 | 5 | 5 | | 3 | 1 | 0 | 5 | 3 | 2 |
| Total Score | | | | | | **17** | **75** | **7** | **20** | **4** | **62** | **37** | **14** |

| ASO | | | | | | W2 w/Ca2+ | | | | | | | |
|---|---|---|---|---|---|---|---|---|---|---|---|---|---|
| Mouse ID | | | | | | ***25*** | ***26*** | ***27*** | ***28*** | ***29*** | ***30*** | ***31*** | ***32*** |
| Death | 75 | | | | | | | | 75 | | | | |
| | Severe | | Moderate | | Mild | | | | | | | | |
| Tonic seizure | 20 | | 15 | | 10 | | | | | | | | |
| Hyperactivity or spasticity | 15 | | 10 | | 5 | | | | | | | | |
| Time required for: | 0.5 h | 1 h | 2 h | 4 h | ≥24 h | | | | | | | | |
| Maintenance of sternal posture | 0 | 4 | 8 | 12 | 20 | 0 | 0 | 0 | 0 | 0 | 0 | 0 | 0 |
| Unstimulated Movement | 0 | 3 | 6 | 9 | 15 | 3 | 3 | 6 | 0 | 3 | 0 | 0 | 0 |
| Movement without Ataxia | 0 | 2 | 4 | 6 | 10 | 6 | 10 | 10 | 0 | 10 | 2 | 2 | 2 |
| Normal Grooming/Eating/Nesting | 0 | 1 | 2 | 3 | 5 | 3 | 2 | 2 | 0 | 1 | 0 | 0 | 0 |
| Total Score | | | | | | **12** | **15** | **18** | **75** | **14** | **2** | **2** | **2** |

| ASO | | | | | | PBS | | | | | | | |
|---|---|---|---|---|---|---|---|---|---|---|---|---|---|
| Mouse ID | | | | | | ***33*** | ***34*** | ***35*** | ***36*** | ***37*** | ***38*** | ***39*** | ***40*** |
| Death | 75 | | | | | 0 | 0 | 0 | 0 | 0 | 0 | 0 | 0 |
| | Severe | | Moderate | | Mild | | | | | | | | |
| Tonic seizure | 20 | | 15 | | 10 | 0 | 0 | 0 | 0 | 0 | 0 | 0 | 0 |
| Hyperactivity or spasticity | 15 | | 10 | | 5 | 0 | 0 | 0 | 0 | 0 | 0 | 0 | 0 |
| Time required for: | 0.5 h | 1 h | 2 h | 4 h | ≥24 h | | | | | | | | |
| Maintenance of sternal posture | 0 | 4 | 8 | 12 | 20 | 0 | 0 | 0 | 0 | 0 | 0 | 0 | 0 |
| Unstimulated Movement | 0 | 3 | 6 | 9 | 15 | 0 | 0 | 0 | 0 | 0 | 0 | 0 | 0 |
| Movement without Ataxia | 0 | 2 | 4 | 6 | 10 | 0 | 0 | 0 | 0 | 0 | 0 | 0 | 2 |
| Normal Grooming/Eating/Nesting | 0 | 1 | 2 | 3 | 5 | 1 | 1 | 0 | 1 | 1 | 1 | 1 | 0 |
| Total Score | | | | | | **1** | **1** | **0** | **1** | **1** | **1** | **1** | **2** |

**Table S5. Raw data from the EvADINT scoring assay for each mouse in Figure 1D (non-targeting ASO).**

| ASO | | | | | | **Full PS** | | | | | |
|---|---|---|---|---|---|---|---|---|---|---|---|
| **Mouse ID** | | | | | | ***135*** | ***136*** | ***137*** | ***138*** | ***139*** | ***140*** |
| **Death** | | | 75 | | | 75 | | | | | |
| | **Severe** | | **Moderate** | | **Mild** | | | | | | |
| **Tonic seizure** | 20 | | 15 | | 10 | | 0 | 0 | 0 | 0 | 0 |
| **Hyperactivity or spasticity** | 15 | | 10 | | 5 | | 10 | 10 | 10 | 10 | 5 |
| **Time required for:** | **0.5 h** | **1 h** | **2 h** | **4 h** | **≥24 h** | | | | | | |
| **Maintenance of sternal posture** | 0 | 4 | 8 | 12 | 20 | | 20 | 20 | 0 | 20 | 0 |
| **Unstimulated Movement** | 0 | 3 | 6 | 9 | 15 | | 15 | 15 | 9 | 15 | 3 |
| **Movement without Ataxia** | 0 | 2 | 4 | 6 | 10 | | 10 | 10 | 6 | 10 | 6 |
| **Normal Grooming/Eating/Nesting** | 0 | 1 | 2 | 3 | 5 | | 5 | 5 | 3 | 5 | 1 |
| **Total Score** | | | | | | **75** | **60** | **60** | **28** | **60** | **15** |

| ASO | | | | | | **Full PS w/Ca2+** | | | | | |
|---|---|---|---|---|---|---|---|---|---|---|---|
| **Mouse ID** | | | | | | ***141*** | ***142*** | ***143*** | ***144*** | ***145*** | ***146*** |
| **Death** | | | 75 | | | | | | | | 75 |
| | **Severe** | | **Moderate** | | **Mild** | | | | | | |
| **Tonic seizure** | 20 | | 15 | | 10 | 0 | 0 | 0 | 0 | 0 | |
| **Hyperactivity or spasticity** | 15 | | 10 | | 5 | 10 | 10 | 10 | 10 | 10 | |
| **Time required for:** | **0.5 h** | **1 h** | **2 h** | **4 h** | **≥24 h** | | | | | | |
| **Maintenance of sternal posture** | 0 | 4 | 8 | 12 | 20 | 12 | 12 | 12 | 12 | 12 | |
| **Unstimulated Movement** | 0 | 3 | 6 | 9 | 15 | 15 | 15 | 9 | 9 | 9 | |
| **Movement without Ataxia** | 0 | 2 | 4 | 6 | 10 | 10 | 10 | 10 | 10 | 10 | |
| **Normal Grooming/Eating/Nesting** | 0 | 1 | 2 | 3 | 5 | 5 | 5 | 5 | 5 | 3 | |
| **Total Score** | | | | | | **52** | **52** | **46** | **46** | **44** | **75** |

| | | | | | | | | | | | |
|---|---|---|---|---|---|---|---|---|---|---|---|
| **ASO** | | | | | | **W2** | | | | | |
| **Mouse ID** | | | | | | ***147*** | ***148*** | ***149*** | ***150*** | ***151*** | ***152*** |
| **Death** | | | 75 | | | | | | | | |
| | **Severe** | | **Moderate** | | **Mild** | | | | | | |
| **Tonic seizure** | 20 | | 15 | | 10 | 0 | 0 | 0 | 0 | 0 | 0 |
| **Hyperactivity or spasticity** | 15 | | 10 | | 5 | 0 | 0 | 0 | 0 | 0 | 0 |
| **Time required for:** | **0.5 h** | **1 h** | **2 h** | **4 h** | **≥24 h** | | | | | | |
| **Maintenance of sternal posture** | 0 | 4 | 8 | 12 | 20 | 0 | 0 | 0 | 0 | 0 | 0 |
| **Unstimulated Movement** | 0 | 3 | 6 | 9 | 15 | 0 | 0 | 0 | 0 | 0 | 0 |
| **Movement without Ataxia** | 0 | 2 | 4 | 6 | 10 | 4 | 4 | 4 | 4 | 4 | 4 |
| **Normal Grooming/Eating/Nesting** | 0 | 1 | 2 | 3 | 5 | 2 | 2 | 2 | 2 | 0 | 1 |
| **Total Score** | | | | | | **6** | **6** | **6** | **6** | **4** | **5** |

| | | | | | | | | | | | |
|---|---|---|---|---|---|---|---|---|---|---|---|
| **ASO** | | | | | | **W2 w/Ca2+** | | | | | |
| **Mouse ID** | | | | | | ***153*** | ***154*** | ***155*** | ***156*** | ***157*** | ***158*** |
| **Death** | | | 75 | | | | | | | | |
| | **Severe** | | **Moderate** | | **Mild** | | | | | | |
| **Tonic seizure** | 20 | | 15 | | 10 | 0 | 0 | 0 | 0 | 0 | 0 |
| **Hyperactivity or spasticity** | 15 | | 10 | | 5 | 0 | 0 | 0 | 0 | 0 | 0 |
| **Time required for:** | **0.5 h** | **1 h** | **2 h** | **4 h** | **≥24 h** | | | | | | |
| **Maintenance of sternal posture** | 0 | 4 | 8 | 12 | 20 | 0 | 0 | 0 | 0 | 0 | 0 |
| **Unstimulated Movement** | 0 | 3 | 6 | 9 | 15 | 0 | 0 | 0 | 0 | 0 | 0 |
| **Movement without Ataxia** | 0 | 2 | 4 | 6 | 10 | 4 | 4 | 4 | 4 | 4 | 4 |
| **Normal Grooming/Eating/Nesting** | 0 | 1 | 2 | 3 | 5 | 0 | 0 | 1 | 2 | 1 | 1 |
| **Total Score** | | | | | | **4** | **4** | **5** | **6** | **5** | **5** |

| | | | | | | | | | | | |
|---|---|---|---|---|---|---|---|---|---|---|---|
| **ASO** | | | | | | **PBS** | | | | | |
| **Mouse ID** | | | | | | ***129*** | ***130*** | ***131*** | ***132*** | ***133*** | ***134*** |
| **Death** | | | 75 | | | | | | | | |
| | **Severe** | | **Moderate** | | **Mild** | | | | | | |
| **Tonic seizure** | 20 | | 15 | | 10 | 0 | 0 | 0 | 0 | 0 | 0 |
| **Hyperactivity or spasticity** | 15 | | 10 | | 5 | 0 | 0 | 0 | 0 | 0 | 0 |
| **Time required for:** | **0.5 h** | **1 h** | **2 h** | **4 h** | **≥24 h** | | | | | | |
| **Maintenance of sternal posture** | 0 | 4 | 8 | 12 | 20 | 0 | 0 | 0 | 0 | 0 | 0 |
| **Unstimulated Movement** | 0 | 3 | 6 | 9 | 15 | 0 | 0 | 0 | 0 | 0 | 0 |
| **Movement without Ataxia** | 0 | 2 | 4 | 6 | 10 | 2 | 2 | 2 | 2 | 2 | 2 |
| **Normal Grooming/Eating/Nesting** | 0 | 1 | 2 | 3 | 5 | 1 | 1 | 1 | 1 | 1 | 1 |
| **Total Score** | | | | | | **3** | **3** | **3** | **3** | **3** | **3** |

**Table S6. Raw data from the EvADINT scoring assay for each mouse in Figure 2 (effect of sugar modifications).**

| ASO | | | | | | Full DNA | | | | | | | |
|---|---|---|---|---|---|---|---|---|---|---|---|---|---|
| Mouse ID | | | | | | *65* | *66* | *67* | *68* | *69* | *70* | *71* | *72* |
| Death | | | 75 | | | 0 | 0 | 0 | 0 | 0 | 0 | 0 | 75 |
| | Severe | | Moderate | | Mild | | | | | | | | |
| Tonic seizure | 20 | | 15 | | 10 | 20 | 0 | 20 | 20 | 20 | 15 | 10 | 0 |
| Hyperactivity or spasticity | 15 | | 10 | | 5 | 0 | 0 | 0 | 0 | 0 | 0 | 0 | 0 |
| Time required for: | 0.5 h | 1 h | 2 h | 4 h | ≥24 h | | | | | | | | |
| Maintenance of sternal posture | 0 | 4 | 8 | 12 | 20 | 20 | 0 | 20 | 20 | 20 | 4 | 4 | 0 |
| Unstimulated Movement | 0 | 3 | 6 | 9 | 15 | 15 | 9 | 15 | 15 | 15 | 6 | 15 | 0 |
| Movement without Ataxia | 0 | 2 | 4 | 6 | 10 | 10 | 10 | 10 | 10 | 10 | 6 | 10 | 0 |
| Normal Grooming/Eating/Nesting | 0 | 1 | 2 | 3 | 5 | 5 | 5 | 5 | 5 | 5 | 5 | 5 | 0 |
| Total Score | | | | | | **70** | **24** | **70** | **70** | **70** | **36** | **44** | **75** |

| ASO | | | | | | Full 2'-O-MOE | | | | | | | |
|---|---|---|---|---|---|---|---|---|---|---|---|---|---|
| Mouse ID | | | | | | *73* | *74* | *75* | *76* | *77* | *78* | *79* | *80* |
| Death | | | 75 | | | 0 | 0 | 0 | 0 | 0 | 0 | 0 | 0 |
| | Severe | | Moderate | | Mild | | | | | | | | |
| Tonic seizure | 20 | | 15 | | 10 | 0 | 0 | 0 | 0 | 0 | 10 | 10 | 0 |
| Hyperactivity or spasticity | 15 | | 10 | | 5 | 10 | 10 | 10 | 10 | 10 | 0 | 0 | 10 |
| Time required for: | 0.5 h | 1 h | 2 h | 4 h | ≥24 h | | | | | | | | |
| Maintenance of sternal posture | 0 | 4 | 8 | 12 | 20 | 4 | 4 | 4 | 4 | 4 | 0 | 0 | 4 |
| Unstimulated Movement | 0 | 3 | 6 | 9 | 15 | 9 | 9 | 9 | 6 | 9 | 0 | 0 | 9 |
| Movement without Ataxia | 0 | 2 | 4 | 6 | 10 | 6 | 6 | 6 | 6 | 6 | 6 | 6 | 6 |
| Normal Grooming/Eating/Nesting | 0 | 1 | 2 | 3 | 5 | 3 | 3 | 3 | 3 | 3 | 3 | 3 | 3 |
| Total Score | | | | | | **32** | **32** | **32** | **29** | **32** | **19** | **19** | **32** |

| ASO | | | | | | Full 2'-O-Me | | | | | | | |
|---|---|---|---|---|---|---|---|---|---|---|---|---|---|
| Mouse ID | | | | | | *81* | *82* | *83* | *84* | *85* | *86* | *87* | *88* |
| Death | | | 75 | | | 0 | 0 | 0 | 0 | 0 | 0 | 0 | 0 |
| | Severe | | Moderate | | Mild | | | | | | | | |
| Tonic seizure | 20 | | 15 | | 10 | 0 | 0 | 0 | 0 | 0 | 0 | 0 | 10 |
| Hyperactivity or spasticity | 15 | | 10 | | 5 | 5 | 10 | 10 | 15 | 10 | 5 | 5 | 0 |
| Time required for: | 0.5 h | 1 h | 2 h | 4 h | ≥24 h | | | | | | | | |
| Maintenance of sternal posture | 0 | 4 | 8 | 12 | 20 | 8 | 4 | 0 | 8 | 4 | 0 | 0 | 0 |
| Unstimulated Movement | 0 | 3 | 6 | 9 | 15 | 6 | 3 | 3 | 9 | 3 | 0 | 3 | 6 |
| Movement without Ataxia | 0 | 2 | 4 | 6 | 10 | 6 | 6 | 4 | 6 | 6 | 6 | 6 | 10 |
| Normal Grooming/Eating/Nesting | 0 | 1 | 2 | 3 | 5 | 3 | 3 | 3 | 3 | 3 | 5 | 3 | 3 |
| Total Score | | | | | | **28** | **26** | **20** | **41** | **26** | **16** | **17** | **29** |

| ASO | | | | | | PBS | | | | | | | |
|---|---|---|---|---|---|---|---|---|---|---|---|---|---|
| Mouse ID | | | | | | *89* | *90* | *91* | *92* | *93* | *94* | *95* | *96* |
| Death | 75 | | | | | 0 | 0 | 0 | 0 | 0 | 0 | 0 | 0 |
| | Severe | | Moderate | | Mild | | | | | | | | |
| Tonic seizure | 20 | | 15 | | 10 | 0 | 0 | 0 | 0 | 0 | 0 | 0 | 0 |
| Hyperactivity or spasticity | 15 | | 10 | | 5 | 0 | 0 | 5 | 0 | 0 | 0 | 0 | 0 |
| Time required for: | 0.5 h | 1 h | 2 h | 4 h | ≥24 h | | | | | | | | |
| Maintenance of sternal posture | 0 | 4 | 8 | 12 | 20 | 0 | 0 | 0 | 0 | 0 | 0 | 0 | 0 |
| Unstimulated Movement | 0 | 3 | 6 | 9 | 15 | 0 | 0 | 0 | 0 | 0 | 0 | 0 | 0 |
| Movement without Ataxia | 0 | 2 | 4 | 6 | 10 | 2 | 2 | 4 | 4 | 4 | 2 | 2 | 4 |
| Normal Grooming/Eating/Nesting | 0 | 1 | 2 | 3 | 5 | 0 | 0 | 1 | 0 | 0 | 0 | 0 | 0 |
| Total Score | | | | | | **2** | **2** | **10** | **4** | **4** | **2** | **2** | **4** |

**Table S7. Raw data from the EvADINT scoring assay for each mouse in Figure 3A.**

For this Figure, the PS-DNA and PBS groups are the same as those used in Figure 2 (the experiments in Figures 2 and 3 were carried out at the same time so those two groups were rightly able to be used for both comparisons). The mouse data for the poly(I:C)-treated mouse is given here:

| ASO | | | | | | Poly (I:C) | | | | | | | |
|---|---|---|---|---|---|---|---|---|---|---|---|---|---|
| Mouse ID | | | | | | *97* | *98* | *99* | *100* | *101* | *102* | *103* | *104* |
| Death | 75 | | | | | 0 | 0 | 0 | 0 | 0 | 0 | 0 | 0 |
| | Severe | | Moderate | | Mild | | | | | | | | |
| Tonic seizure | 20 | | 15 | | 10 | 0 | 0 | 0 | 0 | 0 | 0 | 0 | 0 |
| Hyperactivity or spasticity | 15 | | 10 | | 5 | 0 | 0 | 0 | 0 | 0 | 0 | 0 | 0 |
| Time required for: | 0.5 h | 1 h | 2 h | 4 h | ≥24 h | | | | | | | | |
| Maintenance of sternal posture | 0 | 4 | 8 | 12 | 20 | 0 | 0 | 0 | 0 | 0 | 0 | 0 | 0 |
| Unstimulated Movement | 0 | 3 | 6 | 9 | 15 | 0 | 0 | 0 | 0 | 0 | 0 | 0 | 0 |
| Movement without Ataxia | 0 | 2 | 4 | 6 | 10 | 4 | 4 | 4 | 2 | 2 | 2 | 2 | 2 |
| Normal Grooming/Eating/Nesting | 0 | 1 | 2 | 3 | 5 | 0 | 0 | 0 | 0 | 0 | 0 | 0 | 0 |
| Total Score | | | | | | **4** | **4** | **4** | **2** | **2** | **2** | **2** | **2** |

**Table S8. Raw data from the EvADINT scoring assay for each mouse in Figure 3B.**

| ASO | | | | | | B6 WT / PBS | | | | |
|---|---|---|---|---|---|---|---|---|---|---|
| Mouse ID | | | | | | 159 | 160 | 161 | 162 | 163 |
| Death | 75 | | | | | | | | | |
| | Severe | | Moderate | | Mild | | | | | |
| Seizure | 20 | | 15 | | 10 | | | | | |
| Hyperactivity or spasms | 15 | | 10 | | 5 | | | | | |
| Time required for: | 0.5 h | 1 h | 2 h | 4 h | ≥24 h | | | | | |
| Maintenance of sternal posture | 0 | 4 | 8 | 12 | 20 | 0 | 0 | 0 | 0 | 0 |
| Unstimulated Movement | 0 | 3 | 6 | 9 | 15 | 0 | 0 | 0 | 0 | 0 |
| Movement without Ataxia | 0 | 2 | 4 | 6 | 10 | 4 | 4 | 4 | 4 | 4 |
| Normal Grooming/Eating/Nesting | 0 | 1 | 2 | 3 | 5 | 2 | 2 | 2 | 2 | 2 |
| Total Score | | | | | | 6 | 6 | 6 | 6 | 6 |

| ASO | | | | | | B6 WT / Full DNA | | | | |
|---|---|---|---|---|---|---|---|---|---|---|
| Mouse ID | | | | | | 164 | 165 | 166 | 167 | 168 |
| Death | 75 | | | | | 75 | | 75 | | 75 |
| | Severe | | Moderate | | Mild | | | | | |
| Seizure | 20 | | 15 | | 10 | | | | 10 | |
| Hyperactivity or spasms | 15 | | 10 | | 5 | | 15 | | | |
| Time required for: | 0.5 h | 1 h | 2 h | 4 h | ≥24 h | | | | | |
| Maintenance of sternal posture | 0 | 4 | 8 | 12 | 20 | | 20 | | 20 | |
| Unstimulated Movement | 0 | 3 | 6 | 9 | 15 | | 15 | | 15 | |
| Movement without Ataxia | 0 | 2 | 4 | 6 | 10 | | 10 | | 10 | |
| Normal Grooming/Eating/Nesting | 0 | 1 | 2 | 3 | 5 | | 5 | | 5 | |
| Total Score | | | | | | 75 | 65 | 75 | 60 | 75 |

| ASO | | | | | | MyD88.STING / PBS | | |
|---|---|---|---|---|---|---|---|---|
| Mouse ID | | | | | | 169 | 170 | 171 |
| Death | 75 | | | | | | | |
| | Severe | | Moderate | | Mild | | | |
| Seizure | 20 | | 15 | | 10 | | | |
| Hyperactivity or spasms | 15 | | 10 | | 5 | | | |
| Time required for: | 0.5 h | 1 h | 2 h | 4 h | ≥24 h | | | |
| Maintenance of sternal posture | 0 | 4 | 8 | 12 | 20 | 0 | 0 | 0 |
| Unstimulated Movement | 0 | 3 | 6 | 9 | 15 | 0 | 0 | 0 |
| Movement without Ataxia | 0 | 2 | 4 | 6 | 10 | 4 | 4 | 4 |
| Normal Grooming/Eating/Nesting | 0 | 1 | 2 | 3 | 5 | 1 | 2 | 1 |
| Total Score | | | | | | 5 | 6 | 5 |

| ASO | | | | | | MyD88.Sting / Full DNA | | | |
|---|---|---|---|---|---|---|---|---|---|
| Mouse ID | | | | | | 172 | 173 | 174 | 175 |
| Death | | | 75 | | | 75 | 75 | 75 | |
| | Severe | | Moderate | | Mild | | | | |
| Seizure | 20 | | 15 | | 10 | | | | 10 |
| Hyperactivity or spasms | 15 | | 10 | | 5 | | | | |
| Time required for: | 0.5 h | 1 h | 2 h | 4 h | ≥24 h | | | | |
| Maintenance of sternal posture | 0 | 4 | 8 | 12 | 20 | | | | 20 |
| Unstimulated Movement | 0 | 3 | 6 | 9 | 15 | | | | 15 |
| Movement without Ataxia | 0 | 2 | 4 | 6 | 10 | | | | 10 |
| Normal Grooming/Eating/Nesting | 0 | 1 | 2 | 3 | 5 | | | | 5 |
| Total Score | | | | | | 75 | 75 | 75 | 60 |

**Table S9. Raw data from the EvADINT scoring assay for each mouse in Figure S1A.**

(35 nmol data from Figures 1A)

| ASO | | | | | | *C9orf72* full PS w/Ca2+ (35nmol) | | | | | |
|---|---|---|---|---|---|---|---|---|---|---|---|
| Mouse ID | | | | | | 47 | 48 | 49 | 50 | 51 | 52 |
| Death | | | 75 | | | 0 | 0 | 0 | 0 | 0 | 0 |
| | Severe | | Moderate | | Mild | | | | | | |
| Tonic seizure | 20 | | 15 | | 10 | 0 | 0 | 0 | 0 | 0 | 0 |
| Hyperactivity or spasticity | 15 | | 10 | | 5 | 10 | 0 | 5 | 5 | 0 | 0 |
| Time required for: | 0.5 h | 1 h | 2 h | 4 h | ≥24 h | | | | | | |
| Maintenance of sternal posture | 0 | 4 | 8 | 12 | 20 | 8 | 0 | 12 | 8 | 0 | 12 |
| Unstimulated Movement | 0 | 3 | 6 | 9 | 15 | 9 | 0 | 9 | 9 | 0 | 9 |
| Movement without Ataxia | 0 | 2 | 4 | 6 | 10 | 6 | 4 | 10 | 10 | 2 | 10 |
| Normal Grooming/Eating/Nesting | 0 | 1 | 2 | 3 | 5 | 3 | 3 | 5 | 5 | 0 | 5 |
| Total Score | | | | | | 36 | 7 | 41 | 37 | 2 | 36 |

| ASO | | | | | | *C9orf72* full PS w/Ca2+ (15nmol) | | | | | |
|---|---|---|---|---|---|---|---|---|---|---|---|
| Mouse ID | | | | | | 176 | 177 | 178 | 179 | 180 | 181 |
| Death | | | 75 | | | | | | | | |
| | Severe | | Moderate | | Mild | | | | | | |
| Tonic seizure | 20 | | 15 | | 10 | | | | | | |
| Hyperactivity or spasticity | 15 | | 10 | | 5 | | | | | | |
| Time required for: | 0.5 h | 1 h | 2 h | 4 h | ≥24 h | | | | | | |
| Maintenance of sternal posture | 0 | 4 | 8 | 12 | 20 | 0 | 0 | 0 | 0 | 0 | 0 |
| Unstimulated Movement | 0 | 3 | 6 | 9 | 15 | 0 | 0 | 0 | 0 | 0 | 0 |
| Movement without Ataxia | 0 | 2 | 4 | 6 | 10 | 6 | 6 | 6 | 6 | 6 | 6 |
| Normal Grooming/Eating/Nesting | 0 | 1 | 2 | 3 | 5 | 0 | 1 | 1 | 1 | 0 | 1 |
| Total Score | | | | | | 6 | 7 | 7 | 7 | 6 | 7 |

| ASO | | | | | | *C9orf72* W2 w/Ca2+ (35nmol) | | | | | |
|---|---|---|---|---|---|---|---|---|---|---|---|
| **Mouse ID** | | | | | | **25** | **26** | **27** | **28** | **29** | **30** |
| **Death** | | | 75 | | | 0 | 0 | 0 | 0 | 0 | 0 |
| | **Severe** | | **Moderate** | | **Mild** | | | | | | |
| **Tonic seizure** | 20 | | 15 | | 10 | 0 | 0 | 0 | 0 | 0 | 0 |
| **Hyperactivity or spasticity** | 15 | | 10 | | 5 | 0 | 0 | 0 | 0 | 0 | 0 |
| **Time required for:** | **0.5 h** | **1 h** | **2 h** | **4 h** | **≥24 h** | | | | | | |
| **Maintenance of sternal posture** | 0 | 4 | 8 | 12 | 20 | 0 | 0 | 0 | 0 | 0 | 0 |
| **Unstimulated Movement** | 0 | 3 | 6 | 9 | 15 | 0 | 3 | 3 | 6 | 0 | 0 |
| **Movement without Ataxia** | 0 | 2 | 4 | 6 | 10 | 0 | 6 | 6 | 6 | 6 | 6 |
| **Normal Grooming/Eating/Nesting** | 0 | 1 | 2 | 3 | 5 | 2 | 1 | 2 | 2 | 1 | 1 |
| **Total Score** | | | | | | **2** | **10** | **11** | **14** | **7** | **7** |

| ASO | | | | | | *C9orf72* W2 w/Ca2+ (15nmol) | | | | | |
|---|---|---|---|---|---|---|---|---|---|---|---|
| **Mouse ID** | | | | | | **182** | **183** | **184** | **185** | **186** | **187** |
| **Death** | | | 75 | | | | | | | | |
| | **Severe** | | **Moderate** | | **Mild** | | | | | | |
| **Tonic seizure** | 20 | | 15 | | 10 | | | | | | |
| **Hyperactivity or spasticity** | 15 | | 10 | | 5 | | | | | | |
| **Time required for:** | **0.5 h** | **1 h** | **2 h** | **4 h** | **≥24 h** | | | | | | |
| **Maintenance of sternal posture** | 0 | 4 | 8 | 12 | 20 | 0 | 0 | 0 | 0 | 0 | 0 |
| **Unstimulated Movement** | 0 | 3 | 6 | 9 | 15 | 0 | 0 | 0 | 0 | 0 | 0 |
| **Movement without Ataxia** | 0 | 2 | 4 | 6 | 10 | 6 | 6 | 6 | 6 | 6 | 6 |
| **Normal Grooming/Eating/Nesting** | 0 | 1 | 2 | 3 | 5 | 0 | 0 | 0 | 0 | 0 | 0 |
| **Total Score** | | | | | | **6** | **6** | **6** | **6** | **6** | **6** |

| ASO | | | | | | PBS | | | | | | | |
|---|---|---|---|---|---|---|---|---|---|---|---|---|---|
| **Mouse ID** | | | | | | **33** | **34** | **35** | **36** | **37** | **38** | **39** | **40** |
| **Death** | | | 75 | | | 0 | 0 | 0 | 0 | 0 | 0 | 0 | 0 |
| | **Severe** | | **Moderate** | | **Mild** | | | | | | | | |
| **Tonic seizure** | 20 | | 15 | | 10 | 0 | 0 | 0 | 0 | 0 | 0 | 0 | 0 |
| **Hyperactivity or spasticity** | 15 | | 10 | | 5 | 0 | 0 | 0 | 0 | 0 | 0 | 0 | 0 |
| **Time required for:** | **0.5 h** | **1 h** | **2 h** | **4 h** | **≥24 h** | | | | | | | | |
| **Maintenance of sternal posture** | 0 | 4 | 8 | 12 | 20 | 0 | 0 | 0 | 0 | 0 | 0 | 0 | 0 |
| **Unstimulated Movement** | 0 | 3 | 6 | 9 | 15 | 0 | 0 | 0 | 0 | 0 | 0 | 0 | 0 |
| **Movement without Ataxia** | 0 | 2 | 4 | 6 | 10 | 0 | 0 | 0 | 0 | 0 | 0 | 0 | 2 |
| **Normal Grooming/Eating/Nesting** | 0 | 1 | 2 | 3 | 5 | 1 | 1 | 0 | 1 | 1 | 1 | 1 | 0 |
| **Total Score** | | | | | | **1** | **1** | **0** | **1** | **1** | **1** | **1** | **2** |

**Table S10. Raw data from the EvADINT scoring assay for each mouse in Figure S1B.**

(35 nmol data from Figures 1B)

| **ASO** | | | | | | *Malat1* **Full PS w/Ca2+ (35nmol)** | | | | | |
|---|---|---|---|---|---|---|---|---|---|---|---|
| **Mouse ID** | | | | | | **111** | **112** | **113** | **114** | **115** | **116** |
| **Death** | | | 75 | | | | | | | | |
| | **Severe** | | **Moderate** | | **Mild** | | | | | | |
| **Tonic seizure** | 20 | | 15 | | 10 | 10 | 10 | 10 | 10 | 10 | 10 |
| **Hyperactivity or spasticity** | 15 | | 10 | | 5 | 0 | 0 | 0 | 0 | 0 | 0 |
| **Time required for:** | **0.5 h** | **1 h** | **2 h** | **4 h** | **≥24 h** | | | | | | |
| **Maintenance of sternal posture** | 0 | 4 | 8 | 12 | 20 | 12 | 20 | 0 | 20 | 20 | 20 |
| **Unstimulated Movement** | 0 | 3 | 6 | 9 | 15 | 9 | 15 | 9 | 15 | 15 | 15 |
| **Movement without Ataxia** | 0 | 2 | 4 | 6 | 10 | 10 | 10 | 10 | 10 | 10 | 10 |
| **Normal Grooming/Eating/Nesting** | 0 | 1 | 2 | 3 | 5 | 3 | 5 | 3 | 5 | 5 | 5 |
| **Total Score** | | | | | | **44** | **60** | **32** | **60** | **60** | **60** |

| **ASO** | | | | | | *Malat1* **Full PS w/Ca2+ (15nmol)** | | | | | |
|---|---|---|---|---|---|---|---|---|---|---|---|
| **Mouse ID** | | | | | | **188** | **189** | **190** | **191** | **192** | **193** |
| **Death** | | | 75 | | | | | | | | |
| | **Severe** | | **Moderate** | | **Mild** | | | | | | |
| **Seizure** | 20 | | 15 | | 10 | | 10 | 10 | | 10 | 10 |
| **Hyperactivity or spasms** | 15 | | 10 | | 5 | 10 | | | 10 | | |
| **Time required for:** | **0.5 h** | **1 h** | **2 h** | **4 h** | **≥24 h** | | | | | | |
| **Maintenance of sternal posture** | 0 | 4 | 8 | 12 | 20 | 0 | 8 | 12 | 12 | 12 | 12 |
| **Unstimulated Movement** | 0 | 3 | 6 | 9 | 15 | 0 | 9 | 15 | 15 | 15 | 15 |
| **Movement without Ataxia** | 0 | 2 | 4 | 6 | 10 | 10 | 10 | 10 | 10 | 10 | 10 |
| **Normal Grooming/Eating/Nesting** | 0 | 1 | 2 | 3 | 5 | 2 | 5 | 5 | 5 | 5 | 5 |
| **Total Score** | | | | | | **22** | **42** | **52** | **52** | **52** | **52** |

| **ASO** | | | | | | *Malat1* **W2 w/Ca2+ (35nmol)** | | | | | |
|---|---|---|---|---|---|---|---|---|---|---|---|
| **Mouse ID** | | | | | | **123** | **124** | **125** | **126** | **127** | **128** |
| **Death** | | | 75 | | | | | | | | |
| | **Severe** | | **Moderate** | | **Mild** | | | | | | |
| **Seizure** | 20 | | 15 | | 10 | 0 | 0 | 0 | 0 | 0 | 0 |
| **Hyperactivity or spasms** | 15 | | 10 | | 5 | 5 | 0 | 0 | 0 | 0 | 0 |
| **Time required for:** | **0.5 h** | **1 h** | **2 h** | **4 h** | **≥24 h** | | | | | | |
| **Maintenance of sternal posture** | 0 | 4 | 8 | 12 | 20 | 0 | 0 | 8 | 0 | 0 | 12 |
| **Unstimulated Movement** | 0 | 3 | 6 | 9 | 15 | 9 | 6 | 9 | 6 | 0 | 9 |
| **Movement without Ataxia** | 0 | 2 | 4 | 6 | 10 | 6 | 10 | 10 | 10 | 4 | 10 |
| **Normal Grooming/Eating/Nesting** | 0 | 1 | 2 | 3 | 5 | 3 | 2 | 3 | 3 | 1 | 3 |
| **Total Score** | | | | | | **23** | **18** | **30** | **19** | **5** | **34** |

| ASO | | | | | | *Malat1* W2 w/Ca2+ (15nmol) | | | | | |
|---|---|---|---|---|---|---|---|---|---|---|---|
| Mouse ID | | | | | | 194 | 195 | 196 | 197 | 198 | 199 |
| Death | | | 75 | | | | | | | | |
| | Severe | | Moderate | | Mild | | | | | | |
| Seizure | 20 | | 15 | | 10 | | | | | | |
| Hyperactivity or spasms | 15 | | 10 | | 5 | | | | 5 | | |
| Time required for: | 0.5 h | 1 h | 2 h | 4 h | ≥24 h | | | | | | |
| Maintenance of sternal posture | 0 | 4 | 8 | 12 | 20 | 0 | 0 | 0 | 0 | 0 | 0 |
| Unstimulated Movement | 0 | 3 | 6 | 9 | 15 | 0 | 0 | 0 | 3 | 0 | 0 |
| Movement without Ataxia | 0 | 2 | 4 | 6 | 10 | 6 | 6 | 6 | 6 | 6 | 6 |
| Normal Grooming/Eating/Nesting | 0 | 1 | 2 | 3 | 5 | 0 | 0 | 0 | 2 | 1 | 1 |
| Total Score | | | | | | **6** | **6** | **6** | **16** | **7** | **7** |

| ASO | | | | | | PBS | | | | | |
|---|---|---|---|---|---|---|---|---|---|---|---|
| Mouse ID | | | | | | 200 | 201 | 202 | 203 | 204 | 205 |
| Death | | | 75 | | | | | | | | |
| | Severe | | Moderate | | Mild | | | | | | |
| Seizure | 20 | | 15 | | 10 | | | | | | |
| Hyperactivity or spasms | 15 | | 10 | | 5 | | | | | | |
| Time required for: | 0.5 h | 1 h | 2 h | 4 h | ≥24 h | | | | | | |
| Maintenance of sternal posture | 0 | 4 | 8 | 12 | 20 | 0 | 0 | 0 | 0 | 0 | 0 |
| Unstimulated Movement | 0 | 3 | 6 | 9 | 15 | 0 | 0 | 0 | 0 | 0 | 0 |
| Movement without Ataxia | 0 | 2 | 4 | 6 | 10 | 2 | 2 | 2 | 4 | 2 | 2 |
| Normal Grooming/Eating/Nesting | 0 | 1 | 2 | 3 | 5 | 0 | 0 | 0 | 1 | 0 | 0 |
| Total Score | | | | | | **2** | **2** | **2** | **5** | **2** | **2** |
